# Body coloration and mechanisms of colour production in Archelosauria: The case of deirocheline turtles

**DOI:** 10.1101/556670

**Authors:** Jindřich Brejcha, José Vicente Bataller, Zuzana Bosáková, Jan Geryk, Martina Havlíková, Karel Kleisner, Petr Maršík, Enrique Font

## Abstract

Animal body coloration is a complex trait resulting from the interplay of multiple colour-producing mechanisms. Increasing knowledge of the functional role of animal coloration stresses the need to study the proximate causes of colour production. Here we present a description of colour and colour producing mechanisms in two non-avian archelosaurs, the freshwater turtles *Trachemys scripta* and *Pseudemys concinna*. We compare reflectance spectra; cellular, ultra-, and nano-structure of colour-producing elements; and carotenoid/pteridine derivatives contents in the two species. In addition to xanthophores and melanocytes, we found abundant iridophores which may play a role in integumental colour production. We also found abundant dermal collagen fibres that may serve as thermoprotection but possibly also play role in colour production. The colour of yellow-red skin patches results from an interplay between carotenoids and pteridine derivatives. The two species differ in the distribution of pigment cell types along the dorsoventral head axis, as well as in the diversity of pigments involved in colour production, which may be related to visual signalling. Our results indicate that archelosaurs share some colour production mechanisms with amphibians and lepidosaurs, but also employ novel mechanisms based on the nano-organization of the extracellular protein matrix that they share with mammals.

## 1. Introduction

The evolutionary origins and the selective processes responsible for the maintenance of complex morphological traits remains a persistent challenge in biology. Complex traits may in some cases arise by exaptation of already existing structures into new functions [1]. Animal coloration is one of such complex morphological traits in which composing subunits evolve for various roles and only later are co-opted for signalling or for other functions [2]. Colour-producing elements interact to enhance or reduce each other’s contribution to observable colour [3]. However, colour-producing elements can also develop and evolve independently of each other [4]. Variation in body coloration during evolution is thus the result of two interacting processes: functional integration of colour ornaments and morphological modularity of colour-producing elements [5]. These intriguing relationships between colour-producing subunits and their functional role stress the need to study the proximate causes of colour production [6]. Even though there has been progress in our understanding of colour production and its biological functions [7,8], our knowledge of colour production mechanisms in most groups of animals is still scarce.

Animal colours are produced by ultrastructural elements of the integument interacting with incident light, by light-absorbing pigments, or by a combination of both. In vertebrates, the majority of compounds responsible for the coloration of the integument are found in pigment cells which are derived from the neural crest [9–11]. Pigment cells are classified into five categories: Reflecting dermal pigment cells (iridophores, leucophores), absorbing dermal pigment cells (xanthophores, erythrophores, cyanophores), melanocytes with non-motile organelles (dermal melanocytes), melanocytes with motile organelles (dermal melanophores), and organelle transferring melanocytes (interfollicular and follicular epidermal melanocytes) [12]. Iridophores contain reflecting platelets of guanine, while absorbing dermal pigment cells contain pteridine derivatives in pterinosomes, or carotenoids dissolved in lipids in carotenoid vesicles. Melanocytes and melanophores contain melanins in melanosomes. Single cells with characteristics of two or more of pigment cell types, e.g. xanthophore and iridophore, are occasionally found and are known as mosaic chromatophores [13,14]. Often, pigment cells of the same type are arranged in layers that are roughly parallel to the skin surface. The vertical layering and relative thickness of the different pigment cells, thickness, orientation, number, and geometry of reflecting elements, as well as characteristics of the pigments within the cells determine the final colour of the skin [8,15,16]. Cell types of another embryonic origin, ectodermal keratinocytes of the epidermis and mesenchymal fibroblasts of the dermis, also participate in colour production via reflection of light on the intra/extracellular protein matrix these cells produce [17–21].

Turtles are an early-diverging clade of Archelosauria, the evolutionary lineage of tetrapods leading to crocodiles and birds [22]. Although many turtles have a uniform dull colour, conspicuous striped and spotted patterns are common in all major lineages of turtles (for a comprehensive collection of photographs see [23–26]). These conspicuous colour patterns may be present in the hard-horny skin of shells, and/or in the soft skin of the head, limbs or tail. The dark areas of the skin of turtles may have a threefold origin consisting either of dermal, epidermal, or both epidermal and dermal melanocytes. Colourful bright regions are thought to be the result of the presence of xanthophores in the dermis [27] and their interplay with dermal melanophores [28]. Iridophores have never been shown to play role in coloration of turtles [27,29].

Pigment-bearing xanthophores were first described in the dermis of the Chinese softshell turtle (*Pelodiscus sinensis*) [29]. Xanthophores have also been found sporadically in the dermis of the spiny softshell turtle *Apalone spinifera*, the Murray river turtle (*Emydura macquarii*) and in the painted turtle (*Chrysemys picta*) [27]. Such scarcity of carotenoid/pteridine derivatives-containing cells is in contrast with chemical analyses of the yellow and red regions of the integument of the red-eared slider (*Trachemys scripta elegans*) and *C. picta* [30,31]. Two major classes of carotenoids have been described in the integument of these turtles: short wavelength absorbing apocarotenoids and longer wavelength-absorbing ketocarotenoids [30]. In addition to carotenoids, Steffen et al. [30] speculated that small quantities of some pteridine derivatives could play a minor role in colour production in the integument of turtles. In support of this hypothesis, Alibardi [27] observed pterinosomes with characteristic concentric lamellae in the xanthophores of turtles. However, to this date there is no direct evidence of pteridine derivatives being present in colourful regions of the skin of turtles, or in the skin of any other archelosaur.

Deirochelyinae are freshwater turtles of the family Emydidae, all of which, except one species (the diamond-back terrapin, *Malaclemys terrapin*), display a pattern of conspicuous stripes on the skin of the head and legs [32,33]. This striped pattern usually consists of alternating yellow and black regions, but red regions are also found in some species. The coloration of deirocheline turtles is thought to play a role in mate choice [30,34]. Some species are sexually dichromatic, with males often displaying the brightest, most conspicuous colours (reviewed in [19]); however sexual dichromatism is limited to particular regions of the body, whereas other regions show no sexual differences [35,36]. Colour ornaments are designed to maximize their conspicuousness only in certain, ecologically relevant contexts [37], e.g. when exposed to rivals during contests or to mates during courtship [38,39]. Therefore, it has been suggested that some of the colourful regions found particularly in males may potentially be involved in intersexual communication and mate choice [35,40]. In freshwater turtles courtship takes place underwater and males adopt characteristic positions relative to the female [41]. Deirocheline turtles are characterized by elaborated courtship behaviour including a complex forelimb display known as titillation [41]. Males in most species of deirocheline turtles perform the titillation display facing the female while swimming backwards in front of her, mostly near the surface; however, males of the genus *Pseudemys* titillate swimming above and parallel to the female, facing the same direction as her [41–46]. Thus, depending on the position adopted during courtship (face-to-face vs swim above) males of different species expose different body areas to females. By comparing different deirocheline species, it may be possible to examine whether there is a relationship between colour and colour-producing mechanisms and courtship position in these turtles.

In this study we compare the structure and pigment contents of striped skin in two species of deirocheline freshwater turtles, the pond slider (*Trachemys scripta*) and the river cooter (*Pseudemys concinna*), with contrasting strategies in courtship behaviour. Specifically, we predict that the brightest, most conspicuous skin regions of males will be located precisely in those areas that are most exposed to females during courtship, which will differ depending on the species-specific male position. Males of *T. scripta* court females in a face-to-face position while males of *P. concinna* court females in a swim-above position. In *T. scripta* two of three subspecies (the yellow-bellied slider *T. s. scripta*, and the Cumberland slider *T.s. troosti*) possess only yellow stripes on the head, while the other subspecies (*T.s. elegans*) also has red markings in the postorbital region. All *Pseudemys* turtles show only yellow stripes on the head. We particularly focus on a comprehensive description of the cellular and ultrastructural composition of those body regions that likely play role during courtship behaviour (head, limbs) and on the chemical composition of pigments involved in colour production. We combine multiple approaches to study the mechanisms of colour production: 1) reflectance spectrophotometry to objectively determine the colour of the turtles’ body surfaces, 2) transmission electron microscopy and image analyses to study the composition of the skin and the ultrastructure of colour-producing elements, and 3) liquid chromatography coupled with mass spectrometry to determine the contribution of carotenoid/pteridine derivatives to the turtles’ coloration. We discuss our results from the perspective of the evolution of colour-producing mechanisms in vertebrates. Our results represent the first comprehensive description of colour production in non-avian archelosaurs.

## 2. Materials and Methods

### 2.1 Animals and handling

All turtles examined in this study were collected for purposes of control of invasive freshwater turtles as a continuation of project ‘LIFE-Trachemys’ (LIFE09 NAT/ES000529) to the Servicio de Vida Silvestre, D.G. Medi Natural (Generalitat Valenciana). Turtles (*Pseudemys concinna*: 4 males, 8 females; *Trachemys scripta elegans*: 30 males, 39 females; *Trachemys scripta scripta*: 1 male, 2 females) were captured during 2014-2017 with floating or hoop traps in coastal marshes or artificial ponds at several localities in the Comunidad Valenciana (Spain). Turtles were euthanized with an overdose of sodium pentobarbital (Eutanax, FATRO IBERICA, S.L) by authorized personnel.

### 2.2 Spectral reflectance measurements

To quantify colour, we measured the spectral reflectance of skin between 300 and 700 nm using a spectrophotometer. Reflectance spectra were taken before euthanasia with an Ocean Optics Jaz System (QR400-7-SR-BX: 400 µm reflection probe attached to JAZ-UV-VIS: Deuterium-Tungsten light source module) and a SONY Vaio computer running SpectraSuite software (Ocean Optics, Inc.). The internal trigger was set to 30, integration time to 30, scans average to 30, and boxcar width to 10. The system was calibrated using a Spectralon WS-1 diffuse reflectance standard, also from Ocean Optics. Measurements were taken in a darkened room at a distance of 5 mm and perpendicular to the skin surface (i.e. coincident normal measuring geometry [47]). Turtles were kept alive in clean water inside black plastic tanks and the skin surface was dried with a piece of cloth just before taking reflectance measurements. The main median chin yellow stripe (**CBC**), dorsal head background coloration (**DHC**), main bright/yellow forelimb stripe (**FLBS**) and the postorbital marking on the left side (**PM**) were measured in all specimens of *P. concinna* and *T. s. elegans* (Fig. 1). Unfortunately, as *T. s. scripta* is rarely found in the Valencia region, reflectance spectra could be obtained from only one specimen (male). The yellow zygomatic patch (**YP**) instead of PM was measured in *T. s. scripta* (see Fig. 1B). During postnatal ontogenesis adult males of *T. s. elegans* become melanistic as they get older [48]; therefore, we have not included reflectance spectra of any of the males determined as melanistic in our analyses.

**Figure 1:**
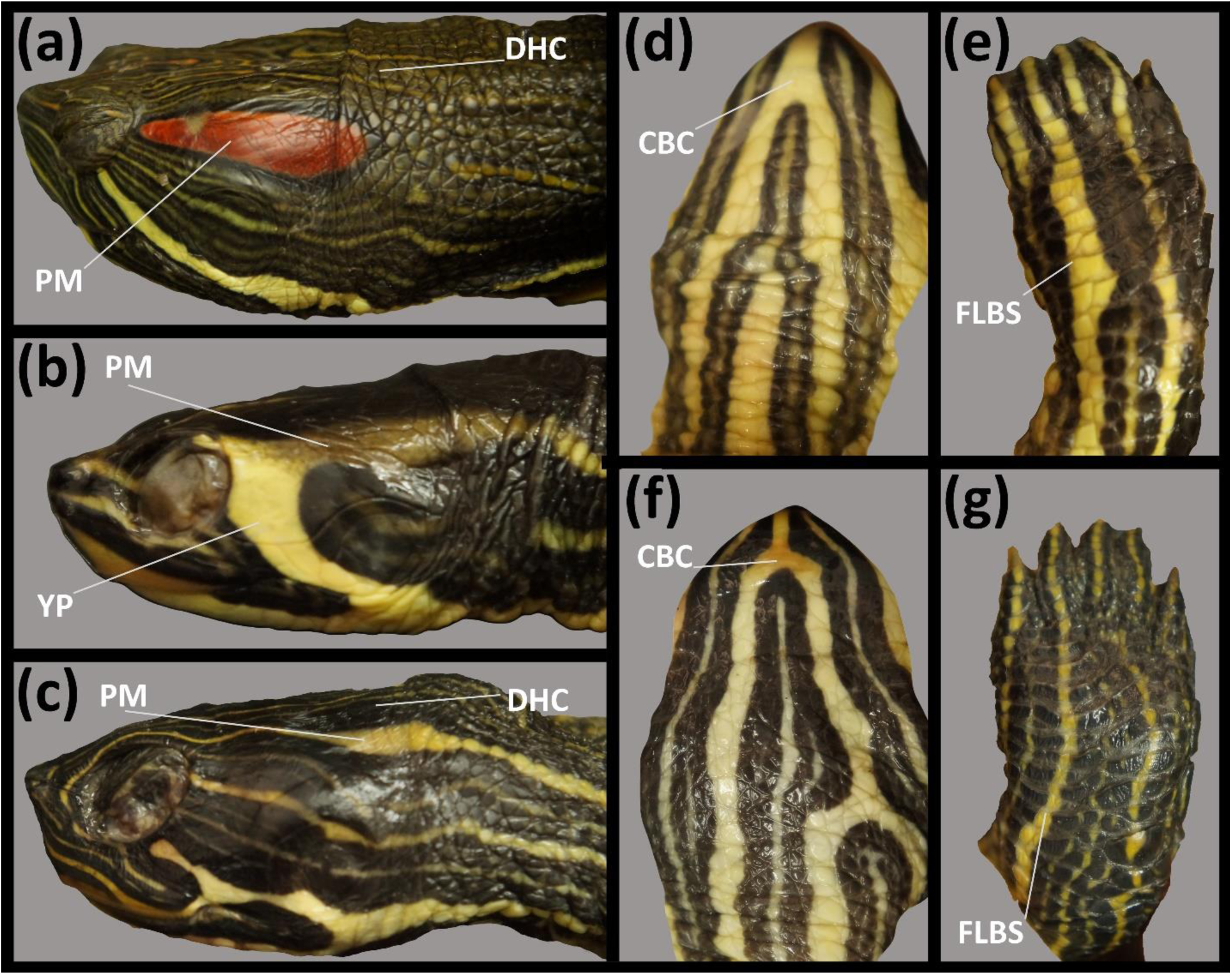
Photographs of skin regions examined in this study. (a) lateral view of the head of *Trachemys scripta elegans*; (b) lateral view of the head of female of *Trachemys scripta scripta*; (c) lateral view of head of female *Pseudemys concinna*; (d) ventral side of head of female of *T. scripta*; (e) front limb of *T. scripta*; (f) ventral side of *P. concinna*; (g) front limb of *P. concinna*. CBC: main median chin yellow stripe; DHC: dorsal head background coloration; FLBS: main bright stripe of the fore limb; PM: postorbital marking; YP: yellow zygomatic patch.

### 2.3 Processing and statistical analyses of reflectance spectra

Reflectance spectra binned by 0.37 nm were restricted to the range between 300 and 700 nm. Calculations were performed in R 3.3.2 [49] using the package PAVO [50]. Spectra were processed by smoothing (span = 0.3) and negative values were fixed to be zero. Mean processed reflectance spectra for each region of each individual of each taxon were summarized by the following variables: luminance or brightness (B1: sum of the relative reflectance over the entire spectral range), hue (H1: wavelength of maximum reflectance), and chroma (relative contribution of a spectral range to total brightness). We calculated five measures of chroma based on the wavelength ranges normally associated with specific colour hues: 300-400 nm (S1.UV), 400-510 nm (S1.blue), 510-605 nm (S1.green), 550-625 nm (S1.yellow), 605-700 nm (S1.red) [50–52]. Reflectance spectra of *T. s. scripta* were not included in the statistical analyses due to the small sample size available for this lineage. Redundancy analysis (RDA) was performed on summary variables scaled to zero mean and unit variance using the package vegan 2.0-10 in R [53]. Species identity was the constraining variable to compare *P. concinna* and *T. s. elegans*. Significance of results was assessed by ANOVA like permutation tests (9,999 permutations) for redundancy analysis (anova.cca) using vegan 2.0-10 package (see script at [Reference to be added]).

### 2.4 Microscopy

To investigate the cellular composition of the turtles’ integument we took samples from CBC, FLBS, and PM regions of *P. concinna* (females = 2) and CBC, DHC, FLBS, and PM regions of *T. s. elegans* (males = 5, females = 3). We also took samples of the CBC, PM, and YP regions of *T. s. scripta* (female = 2). Pieces of integument measuring ca. 1 cm^2^ from target regions were excised immediately after sacrifice and placed in Karnovsky’s fixative (2% paraformaldehyde, 2.5% glutaraldehyde in PB buffer), left overnight at room temperature, and stored in the refrigerator at 4°C for seven days. Smaller pieces (ca. 2 mm^3^) were cut from the original pieces of tissue. These were washed with 0.1M PB, postfixed with 2% osmium tetra oxide (in 0.1M PB solution), dehydrated in an increasing ethanol series, washed with 2% uranyl acetate in 70% ethanol, washed in propylenoxide, after which they were transferred to resin (Durcupan, Sigma). Polymerized resin blocks were cut on a Leica UCT Ultracut ultramicrotome. Semi-thin sections were stained with toluidine blue and observed under bright field and phase contrast in a Nikon Eclipse E800 photomicroscope. Ultra-thin sections were stained with lead citrate and observed and photographed using a FEI Tecnai Spirit G2 TEM equipped with a digital camera (Soft Image System, Morada) and image capture software (TIA 4.7 SP3, FEI Tecnai). Magnification ranged from 1250x to 43000x depending on the structures observed; intensity of the electron beam was adjusted to be in the optimal range for different magnifications.

### 2.5 Reflecting platelets of iridophores

To infer role of iridophores in colour production we analysed the geometric properties of intact reflecting platelets as well as of the empty holes that remain after the reflecting platelets dissolve due to the embedding process. For analyses we used multiple electromicrographs of 4500-9900x magnification: *P. concinna*: PM (N = 8), CBC (N = 8), FLBS (N = 8); *T. s. elegans*: PM (N = 19), CBC (N = 15), FLBS (N = 9). The scale of each image in nm/pixel was set based on the number of pixels in the scale bar of the original electromicrographs. The length of the minor axis (height of reflecting platelet) and of the major axis (width of reflecting platelet) of intact reflecting platelets was measured directly from electromicrographs using the digital callipers utility in ImageJ (1.52a; [54]). Empty holes were analysed automatically by greyscale thresholding and subsequent measurement tools in ImageJ similarly to [16,55]. Briefly, electromicrographs were optimized for contrast and ellipses were fitted inside the platelets’ bodies. Geometric parameters such as length of major axis of ellipse, length of minor axis, and angle between major axis of the ellipse and x axis of electromicrographs were calculated (see electromicrographs of reflecting platelets at [Reference to be added]).

Based on the length of the minor axis of both intact platelets and empty holes we calculated putative reflective wavelengths of reflecting platelets following equation (2) in Morrison [56] as applied by Haisten et al. [57] with refractive index of reflecting platelets 1.83 [58]. However, this model of colour production applies only to reflecting platelets that produce colour through thin-film interference [59], which is characterized by reflecting platelets organized in planes perpendicular to the direction of propagation of incoming light (i.e. parallel to the skin surface). Disorganized reflecting platelets on the other hand act as broadband reflectors producing incoherent scattering and reflect light along the entire wavelength spectrum with little influence on the hue of the specific skin region [16]. Following Saenko et al. [16] we calculated the A/y_0_ ratio to determine the relative amount of reflecting platelets parallel to the skin surface from the Gaussian curve fitted to the distribution of the density of different angles of the ellipses. A is the amplitude of the Gaussian curve above the background of randomly oriented platelets, and y_0_ is the background level of randomly oriented platelets, i. e. intersection of Gaussian curve with y axis. The lower the A/y_0_ ratio the lower the portion of platelets parallel to the skin surface, with 0 corresponding to either no platelets parallel to the skin surface, or completely disorganized platelets [16]. High A and low y_0_ indicate predominant orientation of platelets parallel to the surface and low density of disordered platelets. Full width of half maximum (FWHM) of the peak of Gaussian curve was computed to characterize the spread of the Gaussian curve fitted to the data, i. e. width of angular distribution, as in Saenko et al. [16].

### 2.6 Fourier analysis of spatial distribution of dermal collagen arrays

To determine the role of the abundant extracellular collagen fibres in colour production we performed two dimensional discrete Fourier analyses of the spatial distribution of dermal collagen arrays in the superficial layers of the dermis using a Fourier tool [60] in MATLAB R2018a [61] kindly provided by Dr. Richard Prum (https://prumlab.yale.edu/research/fourier-tool-analysis-coherent-scattering-biological-nanostructures). Because the tool was originally developed for earlier versions of MATLAB, script syntax had to be updated to secure compatibility with current version of MATLAB (see description of changes at [Reference to be added]). The analyses followed published procedures [18,60]. Briefly, from each micrograph (9900x magnification) of cross-sections of the collagen fiber arrays we selected a standardized square portion of the array (800 pixels^2^), and optimized and standardized contrast in Adobe Photoshop (CS3, [62]). The scale of the image in nm/pixel was set based on the number of pixels in the scale bar of the original electromicrographs. The darker pixels of collagen fibres and lighter pixels of mucopolysaccharide were assigned representative refractive indices (1.42 and 1.35 respectively,[63]). We used the Fourier tool with the 2-D fast Fourier transform (FFT2) algorithm [64], resulting in a 2-D Fourier power spectrum expressing the amount of periodicity in the spatial frequencies of the original data. Cumulative analyses were performed using multiple images per each examined region (*P. concinna*: PM, N = 8; CBC, N = 8; FLBS, N = 8; *T. s. elegans*: PM, N = 19; CBC, N = 9; FLBS, N = 9; *T. s. scripta*: CBC, N = 6; YP, N = 11) (see electromicrographs of collagen fibres at [Reference to be added]). For results we pooled images of the CBC region of both *T. scripta* subspecies to increase the sample size for this skin region. Predicted reflectivity in visible spectra of the regions were calculated for 51 concentric bins of the 2-D Fourier power spectra (corresponding to 51 10 nm wide wavelength intervals from 300 to 800 nm) for each region as described in Prum & Torres [60]. To visualize the composite predicted shape of reflectivity from multiple measurements of collagen fibre arrays per region we fitted a smoothed curve (span= 0.3) to the data using the local fitting loess function in R.

### 2.7 Pigment content analyses

Integument from *P. concinna* (N = 3), *T.s. elegans* (N = 2) and *T.s. scripta* (N = 2) was used for carotenoid analyses. Sampled regions were CBC, FLBS, PM, and YP. Small pieces of integument were removed using micro-scissors and tweezers, cleaned mechanically, washed briefly with distilled water to get rid of potential contamination from muscles or body fluids, and frozen at −20°C. Samples were then transported on ice to Prague (Czech Republic) by plane, where they were stored frozen at −20°C until analyses, but never longer than one month. Samples were extracted with 0.5 mL ethyl acetate. The vials containing the extracts were stored for 3 days at room temperature in complete darkness. The extracts were then evaporated to dryness by stream of nitrogen at 27°C and stored at −18°C. Immediately prior to the analyses samples were diluted in 200µl of ethyl acetate (EtOAc).

Standards of astaxanthin, canthaxanthin, lutein, and zeaxanthin were purchased from Sigma Aldrich (Munich, Germany). Stock solutions of the external standards were prepared at a concentration of 0.1 mg/mL by dissolving in EtOAc. The working solution of the mixture of all the studied carotenoids in EtOAc was prepared at concentrations of 50 ng/mL, 100 ng/mL, 200 ng/ml, 500ng/mL and 1000 ng/mL from the stock solutions.

Carotenoids were determined using UPLC system Dionex Ultimate 3000 (Thermo Fischer, USA) coupled with photodiode array (PDA) detector followed by ultra-high resolution accurate mass (HRAM) Q-TOF mass spectrometer (IMPACT II, Bruker Daltonik, Germany). Carotenoid separation was performed on Kinetex C18 RP column (2.6 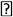 m, 150 x 2.1 mm; Phenomenex, USA) maintained at 35°C using acetonitrile (A), methanol/water 1:1 *v/v* (B) and a mixture of tert-Butyl methyl ether/acetonitrile/methanol - 86:86:8 *v/v/v* (C) as mobile phases for gradient elution (supplementary Table S1) with constant flow rate 0.2 mL/min. Chromatograms were monitored at 445 and 472 nm. The identity of the particular carotenoids was confirmed by HRAM mass spectrometry. Ions were detected in positive mode with electrospray (ESI) as well as atmospheric pressure chemical (APCI) ionization (see details in supplementary Table S2). A chromatogram of the all carotenoid standards mixture in EtOAc at a concentration of 1000 ng/mL is shown as supplementary Fig. S1a, Chromatograms of the individual carotenoid standards with their absorbance spectra are shown in supplementary Fig. S1 c – f.

For analyses of pteridine derivatives we used integument from one individual of each of the examined taxa. Sampled regions were CBC, FLBS, PM, and YP. Samples of integument were removed, treated and transported as for carotenoid analyses. Samples were extracted with dimethyl sulfoxide (DMSO) following a previously published procedure [65].

Standards of 6-biopterin, D-neopterin, leucopterin, pterin, and pterin-6-carboxylic acid (pterin-6-COOH) were purchased from Sigma Aldrich (Munich, Germany). Isoxanthopterin and xanthopterin were obtained from Fluka (Buchs, Switzerland). L-sepiapterin was purchased from the Cayman Chemical Company (Ann Arbor, MI, USA). Erythropterin was kindly provided by Dr. Ron Rutowski. Stock solutions of the external standards were prepared at a concentration of 0.1 mg/mL by dissolving in DMSO. The working solution of the mixture of all the studied pteridine derivatives in DMSO was prepared at a concentration of 0.01 mg/ml from the stock solutions.

All chromatographic measurements were carried out in a HPLC system Agilent series 1290 coupled with a Triple Quad 6460 tandem mass spectrometer (Agilent Technologies, Waldbronn, Germany). For data acquisition, the Mass Hunter Workstation software was used. A ZIC^®^-HILIC (4.6 mm × 150 mm, 3.5 μm) column, based on zwitterionic sulfobetaine groups, was chosen (Merck, Darmstadt, Germany). The chromatographic conditions were adapted from Kozlík et al. [66]. The isocratic elution at a flow rate of 0.5 ml/min with the mobile phase consisted of acetonitrile/5mM ammonium acetate, pH 6.80 at a volume ratio of 85:15 (*v/v*) was used for the separation. Tandem mass spectrometric (MS/MS) measurements were performed in the selected reaction monitoring mode (SRM) with positive ionization. SRM conditions used for MS/MS determination are listed in supplementary Table (S3). Under these conditions pteridine derivatives eluted with the following retention times (min): L-sepiapterin *t* = 7.00, pterin *t* = 9.20, isoxanthopterin *t* = 12.5, 6-biopterin *t* = 12.6, xantopterin *t* = 16.4, leukopterin *t* = 18.4, erythropterin *t* = 25.0, pterin-6-carboxylic acid *t* = 28.0 and D-neopterin *t* = 28.8. A chromatogram of the mixture of all pteridine derivative standards in DMSO at a concentration of 0.01 mg/mL, measured in the TIC (total ion current) mode is shown as supplementary Fig. S2a, SRM chromatograms of the individual derivatives are shown as supplementary Fig. S2b.

## 3. Results

### 3.1 Spectral reflectance

Average standardized reflectance spectra of *Pseudemys concinna* and *Trachemys scripta* are shown in Fig. 2 (see reflectance data of each individual at [Reference to be added]). The dorsal head background coloration (DHC) of both *T. scripta* and *P. concinna* has a flat reflectance spectrum and low overall reflectance. Moreover, the reflectance of DHC is further reduced beyond 550 nm. The red postorbital marking (PM) of *T. s. elegans* also has a relatively flat reflectance spectrum, with low overall reflectance but, in contrast to DHC, the red PM of *T. s. elegans* shows increased reflectance at wavelengths longer than 550 nm giving rise to a plateau between 600-700 nm. All reflectance spectra from yellow regions (i. e. main median chin stripe – CBC, and main forelimb stripe – FLBS, of both species, and PM of *P. concinna* as well as zygomatic patch – YP of *T. s. scripta*) are similar in that they show two peaks, one minor peak in the UV part of the spectrum between 300 and 400 nm and a larger, primary peak between 500 and 600 nm. The width, shape and height of the primary peak differ slightly among the yellow regions.

**Figure 2:**
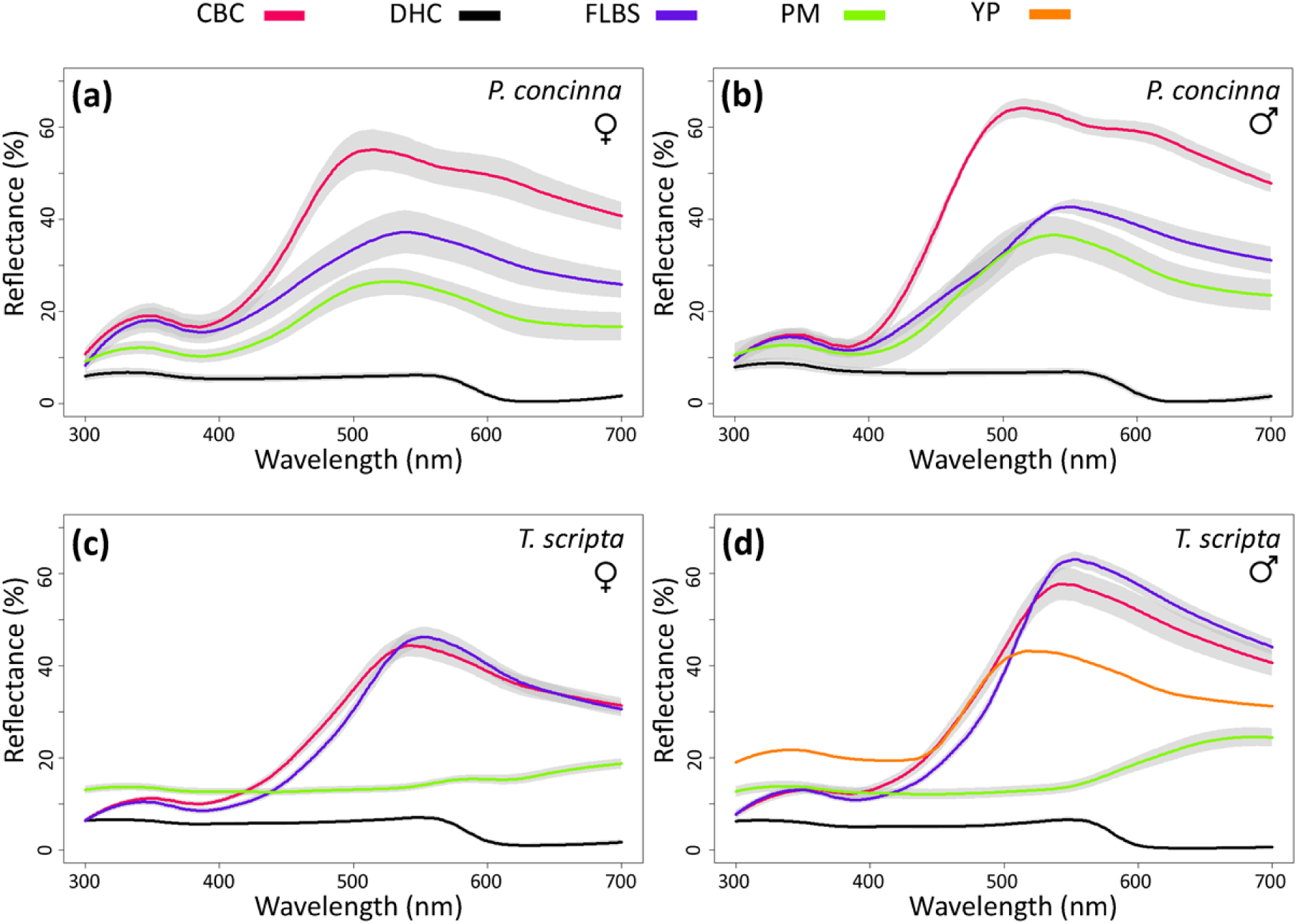
Mean (+/-SEM) reflectance spectra of focal regions of (a) females of *Pseudemys concinna* (N=8), (b) males of *P. concinna* (N=4), (c) females of *Trachemys scripta elegans* (N=39), and (b) males of *T. s. scripta elegans* (N=30). CBC: main median chin yellow stripe; DHC: dorsal head background coloration; FLBS: main bright stripe of the fore limb; PM: postorbital marking. The yellow zygomatic patch (YP) of *T. s. scripta* (N=1) is shown in (d).

Means of summary variables derived from reflectance spectra and their standard deviations for each region of each taxon are shown in Table 1 (see summary variables of each individual at [Reference to be added]). Summary variables derived from reflectance spectra of one individual *T. s. scripta* overlapped with summary variables (mean +/-standard deviation) of *T. s. elegans* but fall outside the values (mean +/-standard deviation) of the summary variables for *P. concinna.*

**Table 1:** Summary colour variables for each taxon (*Pseudemys concinna, Trachemys scripta elegans, Trachemys scripta scripta*) involved in this study. B1 – sum of the relative reflectance over the entire spectral range; H1 – wavelength of maximum reflectance; relative contributions of a spectral ranges to the total brightness: 300-400nm (S1.UV), 400-510nm (S1.blue), 510nm-605nm (S1.green), 550nm-625nm (S1.yellow), 605-700nm (S1.red). CBC: main median chin yellow stripe; DHC: dorsal head background coloration; FLBS: main bright stripe of the fore limb; PM: postorbital marking; YP: yellow zygomatic patch. Note that for *T. s. scripta* the values of summary variables resulted from measurements of only one specimen.

| Taxon (number of individuals) | Region | Variable (SD) |  |  |  |  |  |  |
| --- | --- | --- | --- | --- | --- | --- | --- | --- |
|  |  | B1 | S1.UV | S1.blue | S1.green | S1.yellow | S1.red | H1 |
| *P. concinna* (N=12) | CBC | 15584(2670) | 0.102(0.034) | 0.272(0.019) | 0.34(0.023) | 0.26(0.019) | 0.294(0.025) | 515.25(10.244) |
|  | DHC | 1904.4282(713.37) | 0.357(0.046) | 0.348(0.013) | 0.272(0.037) | 0.141(0.024) | 0.03(0.035) | 365.167(85.58) |
|  | FLBS | 10572.43(3027.39) | 0.14(0.037) | 0.256(0.03) | 0.338(0.029) | 0.257(0.026) | 0.274(0.031) | 540.167(12.511) |
|  | PM | 7775.415(2765.69) | 0.147(0.03) | 0.271(0.028) | 0.347(0.029) | 0.248(0.022) | 0.242(0.037) | 529.75(11.787) |
| *T. s. elegans* (N=68) | CBC | 11923.4(4287.2) | 0.095(0.036) | 0.221(0.027) | 0.382(0.036) | 0.291(0.027) | 0.31(0.029) | 547.882(27.173) |
|  | DHC | 1792.31(1070) | 0.353(0.042) | 0.354(0.027) | 0.28(0.029) | 0.134(0.031) | 0.02(0.053) | 432.088(119.64) |
|  | FLBS | 12026.9(3134.57) | 0.086(0.025) | 0.186(0.027) | 0.406(0.024) | 0.318(0.023) | 0.33(0.029) | 552.912(10.215) |
|  | PM | 5991.65(2426.26) | 0.223(0.037) | 0.231(0.045) | 0.237(0.029) | 0.207(0.032) | 0.316(0.089) | 622.382(123.418) |
| *T. s. scripta* (N=1) | CBC | 11467.67 | 0.094 | 0.229 | 0.375 | 0.28 | 0.309 | 518 |
|  | DHC | 801.896 | 0.29 | 0.37 | 0.346 | 0.159 | 0 | 549 |
|  | FLBS | 10980.87 | 0.078 | 0.192 | 0.405 | 0.309 | 0.334 | 541 |
|  | YP | 12102.164 | 0.172 | 0.252 | 0.323 | 0.239 | 0.261 | 518 |

The ordination plot resulting from redundancy analyses (RDA) based on summary variables of both *T. s elegans* and *P. concinna* is shown in Fig. 3 a. There are significant differences in coloration between *P. concinna* and *T. s. elegans* (ANOVA-like permutation test, F = 16.39, p < 0.001). Constrained axis (RDA1) explains 17% of the total variance in the data. The first residual axis (PC1) explains 18% and the second residual axis (PC2) explains 17% of the total variance. Even though our primary goal was not to investigate sexual dichromatism, the results suggest sexual differences in the reflectance of the red and yellow skin patches in *T. scripta* and, to a lesser extent, *P. concinna* (Fig. 2, See the supplementary materials for details).

**Figure 3:**
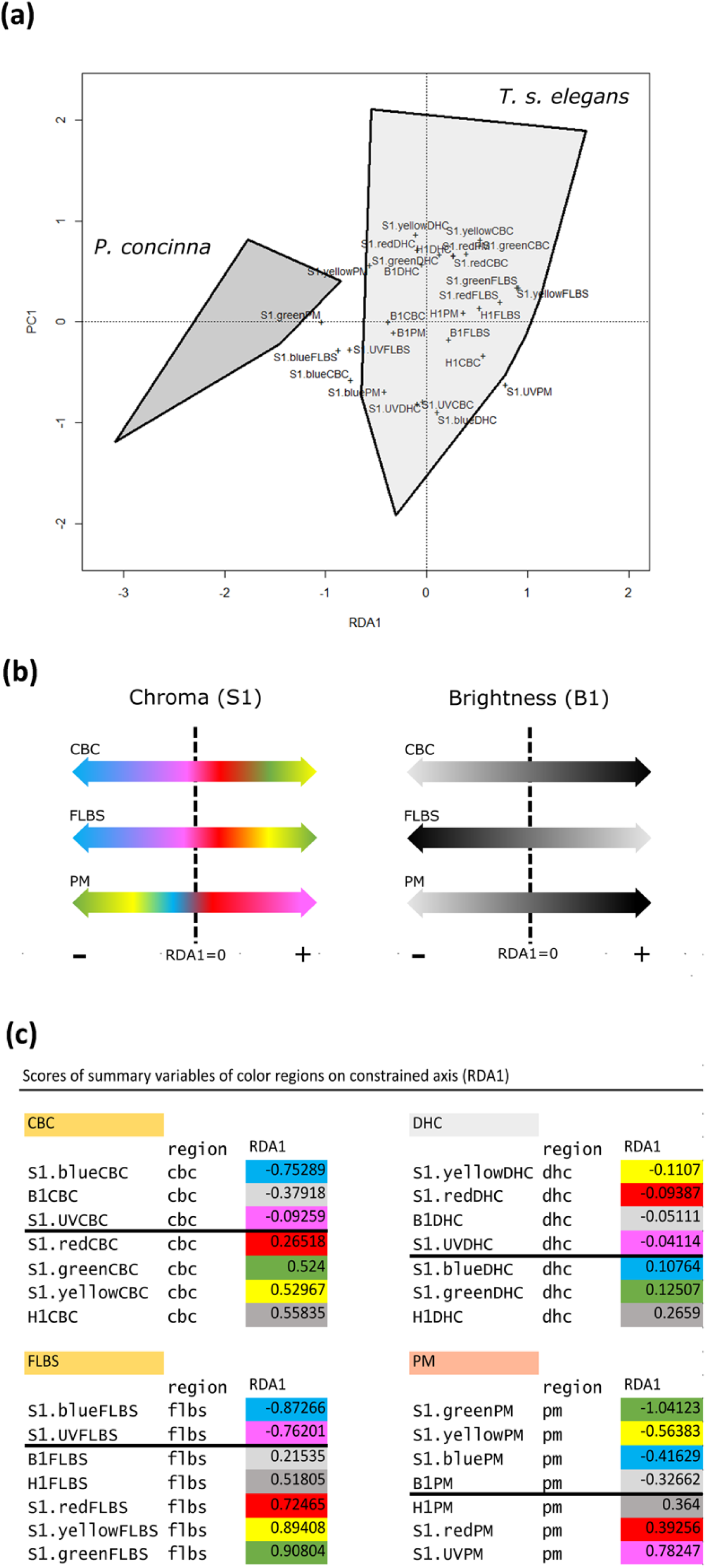
Overall differences in colour between the two studied species. (a) Biplot representing results of RDA of summary variables derived from reflectance spectra of *Pseudemys concinna* and *Trachemys scripta elegans*. First axis (RDA1) is constrained by species (explains 17% of total variance). The first residual axis (PC1) explains 18% of total variance. B1: sum of the relative reflectance over the entire spectral range; H1: wavelength of maximum reflectance; relative contributions of a spectral ranges to the total brightness: 300-400 nm (S1.UV), 400-510 nm (S1.blue), 510-605 nm (S1.green), 550-625 nm (S1.yellow), 605-700 nm (S1.red). CBC: main median chin yellow stripe; DHC: dorsal head background coloration; FLBS: main bright stripe of the fore limb; PM: postorbital marking; YP: yellow zygomatic patch. (b) Schema of the differences between species (distribution along RDA1 axis) in chroma (S1) and brightness (B1) of examined regions. Note that the DHC region aligned rather more with the first residual axis (PC1). (c) Table of scores of summary variables of all colour regions of both species on RDA1 axis. Thick line denotes zero.

Variables pertaining to background colour (DHC) are associated with PC1 rather than with RDA1 (compare values of RDA1 scores of the DHC to scores of summary variables of other regions – Fig. 3B) and contribute very little to colour differences between both species. Interpretation of the RDA1 axis, and therefore of differences between both species, seems to be region specific. *Pseudemys concinna* shows increased brightness (B1) along the RDA1 axis in CBC and PM, but decreased brightness in FLBS compared to *T. s. elegans*. The yellow CBC and FLBS of *P. concinna* show higher chroma (S1) in the blue and UV segments of reflectance spectra, but the yellow CBC and FLBS *T. s.* elegans have higher chroma (S1) in the yellow, red and green segments. The yellow PM of *P. concinna* shows higher chroma in the green, yellow and blue segments of reflectance spectra, while the red PM of *T. s. elegans* shows higher chroma in the red segment, but also in the UV segment of reflectance spectra (Fig. 3 b). The increased chroma in the UV segment of the PM in *T. s. elegans* is a consequence of the overall low reflectance of this region rather than exceptionally high reflectance in the UV. Reported patterns of interspecific differences did not change when the PM region was not included in the analyses (data not shown here but see R script together with input data at [Reference to be added]).

In short, the major differences in colour of examined regions between both species examined are: 1) shorter wavelengths contribute more to yellow colour of the CBC and FLBS of *P. concinna* compared to *T. s. elegans*; 2) CBC and PM are brighter in *P. concinna*, but the FLBS is brighter in *T. s. elegans*.

### 3.2 Microscopy

The epidermis is separated from the dermis by a conspicuous basal lamina (solid arrows in Figs. 4, 5). The epidermis differs in thickness between regions and contains multiple layers of keratinocytes. The corneous layer of the epidermis is relatively thin in all regions except in FLBS where it represents about one fifth of the thickness of the entire epidermis (of all the regions examined only the FLBS is covered by scales). Arrays of collagen fibres are highly abundant in the dermis of every region examined. Four basic pigment cells types are present in the integument of all studied turtles: epidermal melanocytes, and dermal iridophores, melanocytes, and xanthophores (Figs. 4; 5; 6). We have not observed finger-like projections of dermal melanocytes, that are characteristic for dermal melanophores with motile organelles [12]. Mosaic pigment cells were also found in the dermis of *T. s. elegans* (Fig. 6 d, g, h). The pigment cell types present in each of the studied regions are summarized in Table 2.

**Table 2:** Contribution (✔ present/ ✖ absent) of different pigment cell types in particular regions of integument in studied taxa (*Pseudemys concinna, Trachemys scripta elegans, Trachemys scripta scripta*). CBC: main median chin yellow stripe; DHC: dorsal head background coloration; FLBS: main bright stripe of the fore limb; PM: postorbital marking; YP: yellow zygomatic patch.

| Taxon | Region | Pigment cell type |  |  |  |
| --- | --- | --- | --- | --- | --- |
|  |  | epidermal<br>melanocytes | iridophores | dermal<br>melanocytes | xantophores |
| *P. concinna* | CBC | × | ✓ | × | ✓ |
|  | DHC | ✓ | ✓ | ✓ | ✓ |
|  | FLBS | × | ✓ | × | ✓ |
|  | PM | × | ✓ | × | ✓ |
| *T. s. elegans* | CBC | × | ✓ | × | ✓ |
|  | DHC | ✓ | ✓ | ✓ | ✓ |
|  | FLBS | × | ✓ | × | ✓ |
|  | PM | ✓ | × | × | ✓ |
| *T. s. scripta* | CBC | × | ✓ | × | ✓ |
|  | DHC | ✓ | ✓ | ✓ | ✓ |
|  | FLBS | × | ✓ | × | ✓ |
|  | YP | × | × | × | ✓ |

**Figure 4:**
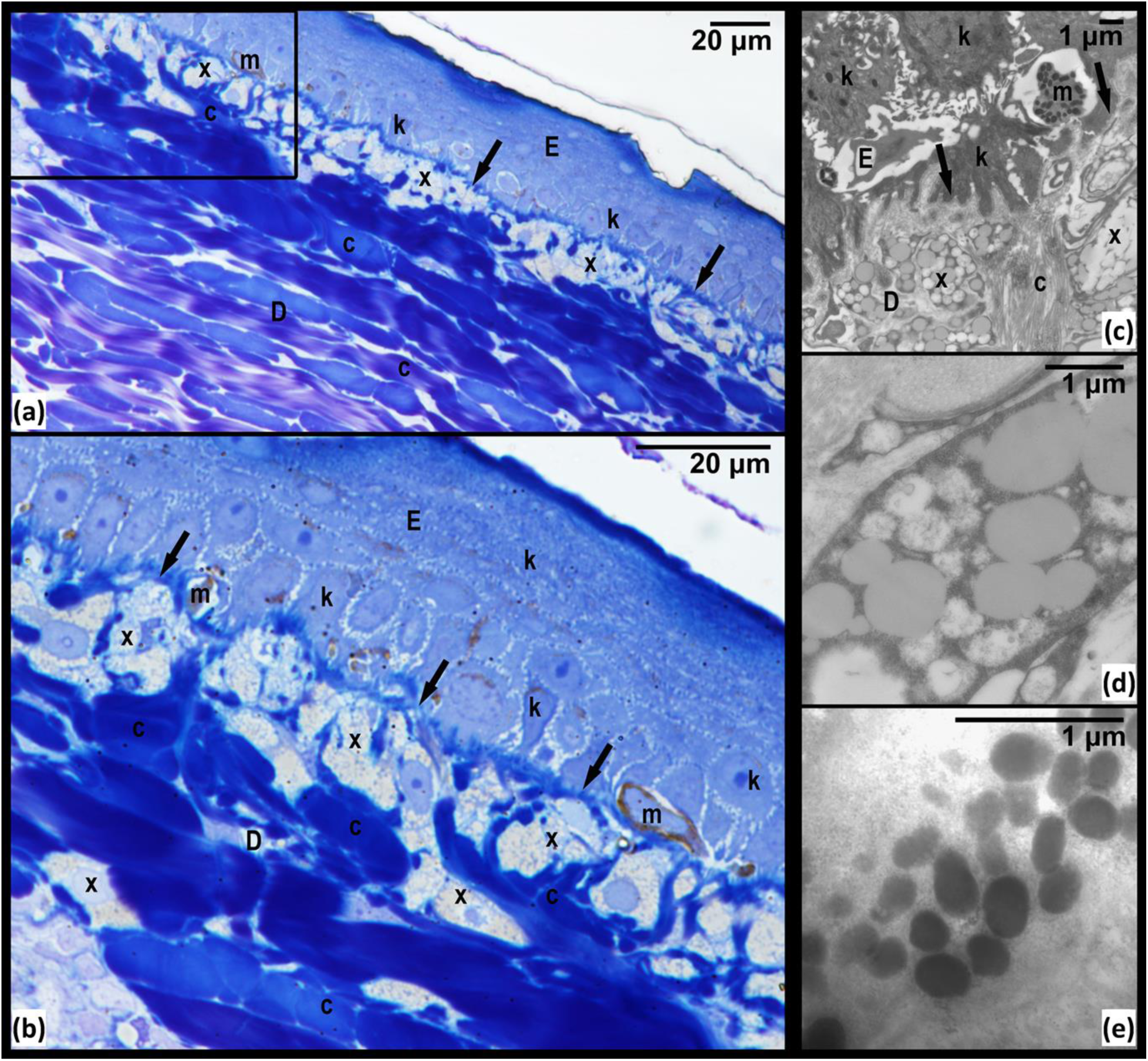
Red postorbital region (PM) of *Trachemys scripta elegans*. (a) micrograph of semi-thin section stained with toluidine blue viewed under light microscope; (b) area denoted by rectangle in (a) viewed at higher magnification; (c) electromicrograph showing xanthophores residing in dermis under basal lamina (solid arrows); (d) closer view of xanthophore and its carotenoids vesicles; (e) detail of melanosomes of epidermal melanocyte. D: dermis; E: epidermis; c: collagen fibres; k: keratinocyte; m: melanocyte; x: xanthophore; arrows: basal lamina.

**Figure 5:**
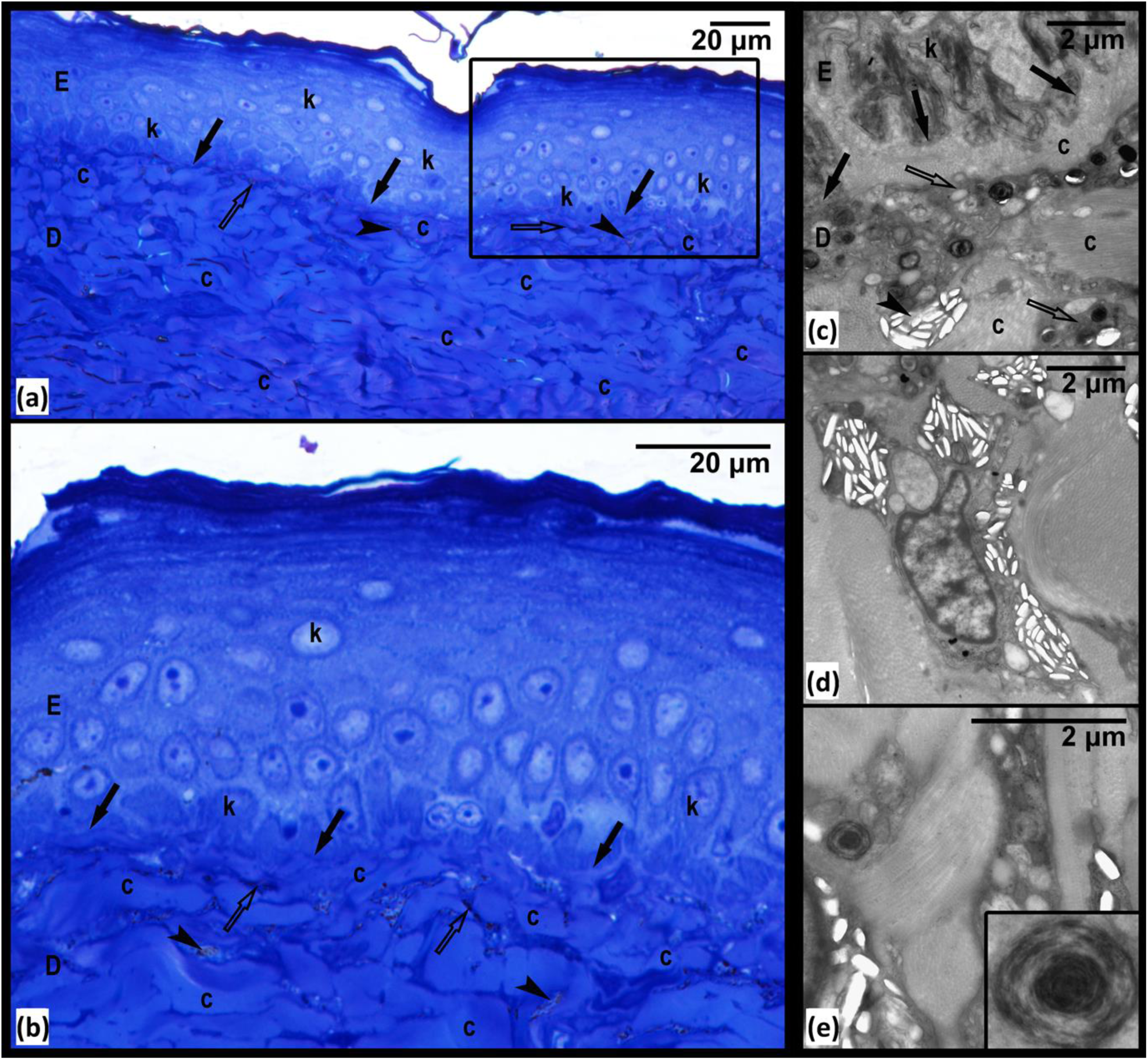
Yellow postorbital region (PM) of *Pseudemys concinna*. (a) semi-thin section stained by toluidine blue viewed under light microscope; (b) area denoted by rectangle in (a) viewed at higher magnification; (c) Electromicrograph showing xanthophores and iridophores in the dermis; (d) closer view of iridophore; (e) closer look on xanthophore with detail of pterinosome in the inset. D: dermis; E: epidermis; c: collagen fibres; k: keratinocyte; solid arrows: basal lamina; clear arrows: xanthophore; arrowheads: iridophores.

**Figure 6:**
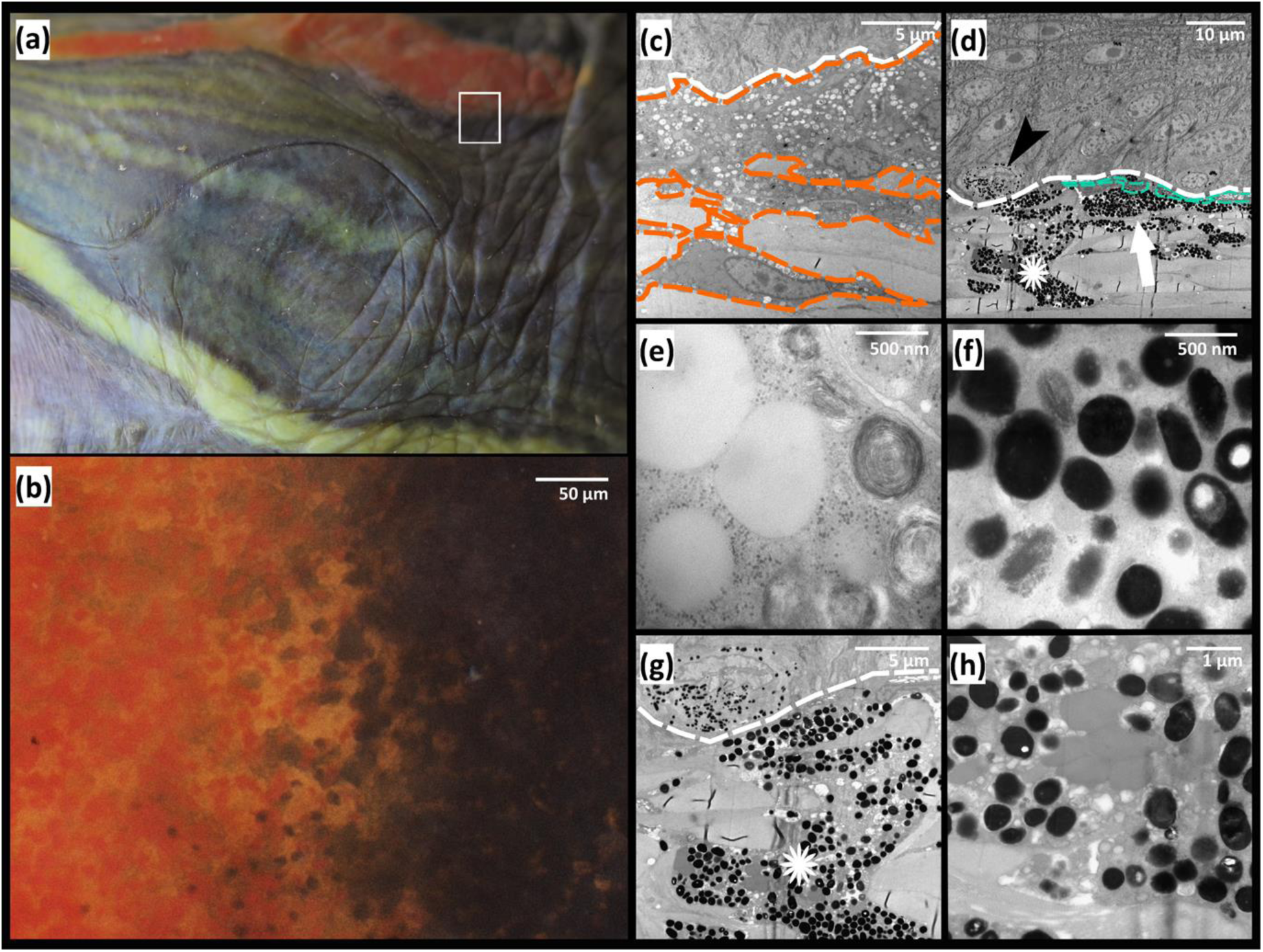
Boundary between the red postorbital region (PM) and its surrounding black line of *Trachemys scripta elegans*; (a) photography of the lateral view of the head, region of interest highlighted by white rectangle. (b) edge between two different regions viewed under binocular microscope; (c) electromicrograph of section of red postorbital region, orange line: xanthophores; white line: basal lamina; (d) edge of black region, turquoise line: iridophore; white line: basal lamina; black arrowhead: epidermal melanocyte; white arrow: dermal melanocytes; asterisk: mosaic chromatophore; (e) detail of vesicles of xanthophores of the red postorbital region – carotenoid vesicles and pterinosomes; (f) detail of vesicles of dermal melanocytes of black region, melanosomes; (g) detailed view of mosaic chromatophore in the dermis. White line: basal lamina; asterisk: mosaic chromatophore; (h) high magnification of vesicles of mosaic chromatophore, showing co-localization of carotenoid vesicles and melanosomes.

Epidermal melanocytes are present in the epidermis in the PM and DHC of *T. scripta* (Figs. 7 e; 8 a,b,c), and DHC of *P. conncina*. Epidermal melanocytes are located just above the basal lamina and are surrounded by keratinocytes. In samples with epidermal melanocytes the surrounding keratinocytes contain melanosomes. In some samples, however, the epidermal melanocytes are contracted and contain only a few melanosomes; those samples lack melanosomes deposited in keratinocytes. The PM of *T. s. elegans* contains mostly contracted epidermal melanocytes and almost no transferred melanosomes (Fig. 8 a). In contrast, the PM of *T. s. scripta* contains enlarged epidermal melanocytes and melanosomes distributed amongst many keratinocytes (Fig. 8 b, c). *Pseudemys concinna* does not have epidermal melanocytes in the PM (Fig. 5, Fig. 8 e, f).

**Figure 7:**
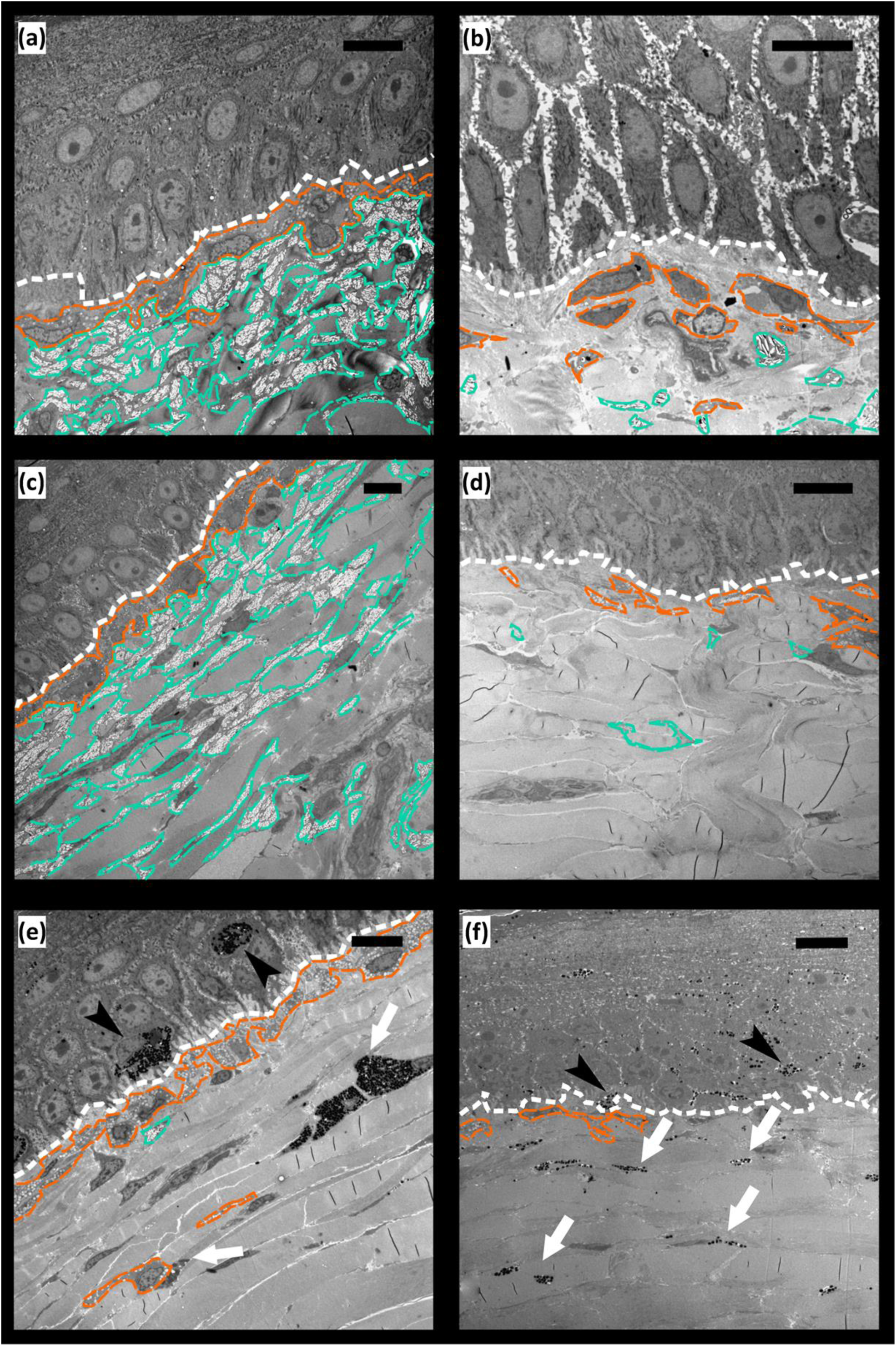
Electromicrographs of different regions of turtles’ integument. (a) yellow chin (CBC) region of *Trachemys scripta*; (b) CBC region of *Pseudemys concinna*; (c) yellow front limb (FLBS) region of *T. scripta*; (d) FLBS region of *P. concinna*; (e) Dorsal dark head (DHC) region of *T. s. elegans*; (f) DHC region of *T. s. scripta*. White line: basal lamina; orange line: xanthophores; turquoise line: iridophores; black arrowheads: epidermal melanocytes; white arrows: dermal melanocytes. Scale bar represents 10µm. Note that magnification differs.

**Figure 8:**
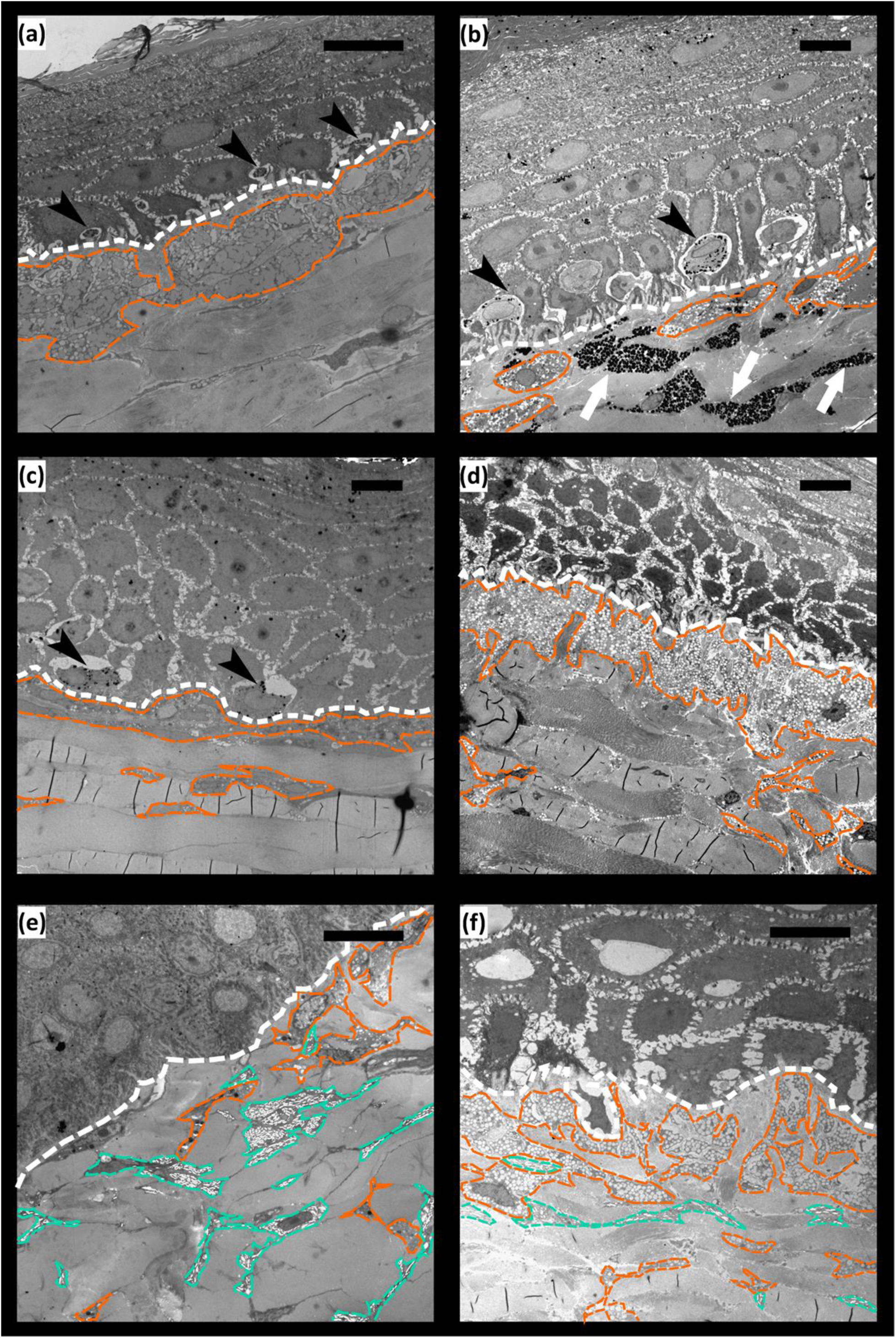
Electromicrographs of different regions of turtles’ integument. (a) red postorbital (PM) region of *Trachemys scripta elegans*, note small epidermal melanocytes and no melanosomes in keratinocytes; (b) red PM region of melanistic male of *T. s. elegans*, note enlarged epidermal melanocytes, but few melanosomes in keratinocytes; (c) dark PM region of *T.s. scripta*, note enlarged epidermal melanocytes and abundant melanosomes in keratinocytes; (d) yellow zygomatic patch (YP) region of *T.s. scripta*, note that there are no epidermal melanocytes; (e) yellow PM region of *P. concinna*; (f) yellow PM region of different specimen of *P. concinna* than in e), see the difference in abundance of xanthophores and iridophores, here xanthophores form almost continuous layer, it is possible that these differences are due to the different localization within the PM stripe on head, or there these differences may be due to intraspecific variability. White line: basal lamina; orange line: xanthophores; turquoise line: iridophores; black arrowheads: epidermal melanocytes; white arrows: dermal melanocytes. Scale bar represents 10µm. Note that magnification differs.

Iridophores are present especially in the dermis of the CBC, FLBS, and PM of *P. concinna*, and in the CBC, FLBS of *T. scripta*, but are missing in the PM and YP of *T. scripta*. Iridophores are present also in the DHC, but these are very rare, small and seem haphazardly distributed (Fig 7 e). The iridophores of *P. concinna* do not form a continuous layer. In the CBC and FLBS of *P. concinna* iridophores are rather scarce (Fig. 7 b, d), but in the PM of *P. concinna* they are more abundant (Fig. 8 e). In the CBC and FLBS of *T. scripta* iridophores are abundant and form an almost continuous layer several cells thick (Fig. 7 a, c).

Dermal melanocytes are abundant in the DHC (Fig. 7 e, f) and in the narrow black lines outlining the yellow-red regions (Fig. 6). In the DHC melanocytes are dispersed throughout the upper parts of dermis but do not generally form a continuous layer. In contrast, the dermal melanocytes in the narrow black lines form a continuous layer. Dermal melanocytes were present also in the PM of a melanistic male of *T. s. elegans* (Fig. 8 b).

Xanthophores are present in the dermis of all regions examined just beneath the basal lamina and just above iridophores when those are present. In the CBC, FLBS (Fig. 7 b,d) and PM (Fig.8 e, f) of *P. concinna* the xanthophores are sparse and do not form a continuous layer. The xanthophores form a thick continuous layer in the PM of *T. s. elegans* (Fig. 8 a) and in the YP of *T. s. scripta* (Fig. 8 d). In the CBC and FLBS of *T. s. elegans* (Fig. 7 a, c) and PM of *T. s. scripta* (Fig. 8 c) the xantophore layer is also continuous but its thickness is reduced compared to the PM of *T. s. elegans* or YP of *T. s. scripta*. Xanthophores are also found in the DHC where they form aggregates of interconnected cells, but not a continuous layer (Fig 7 e). Xanthophores contain three types of pigment bearing organelles: two types of carotenoid vesicles with different electron density but without internal structure, and pterinosomes (Fig. 9). Unlike carotenoid vesicles, mature pterinosomes are characterized by the presence of concentric lamellae [67,68]. Mosaic pigment cells containing carotenoid vesicles, pterinosomes and melanosomes are present at the boundary between the PM and its black surrounding line in *T. s. elegans* (Fig. 6 g, h).

**Figure 9:**
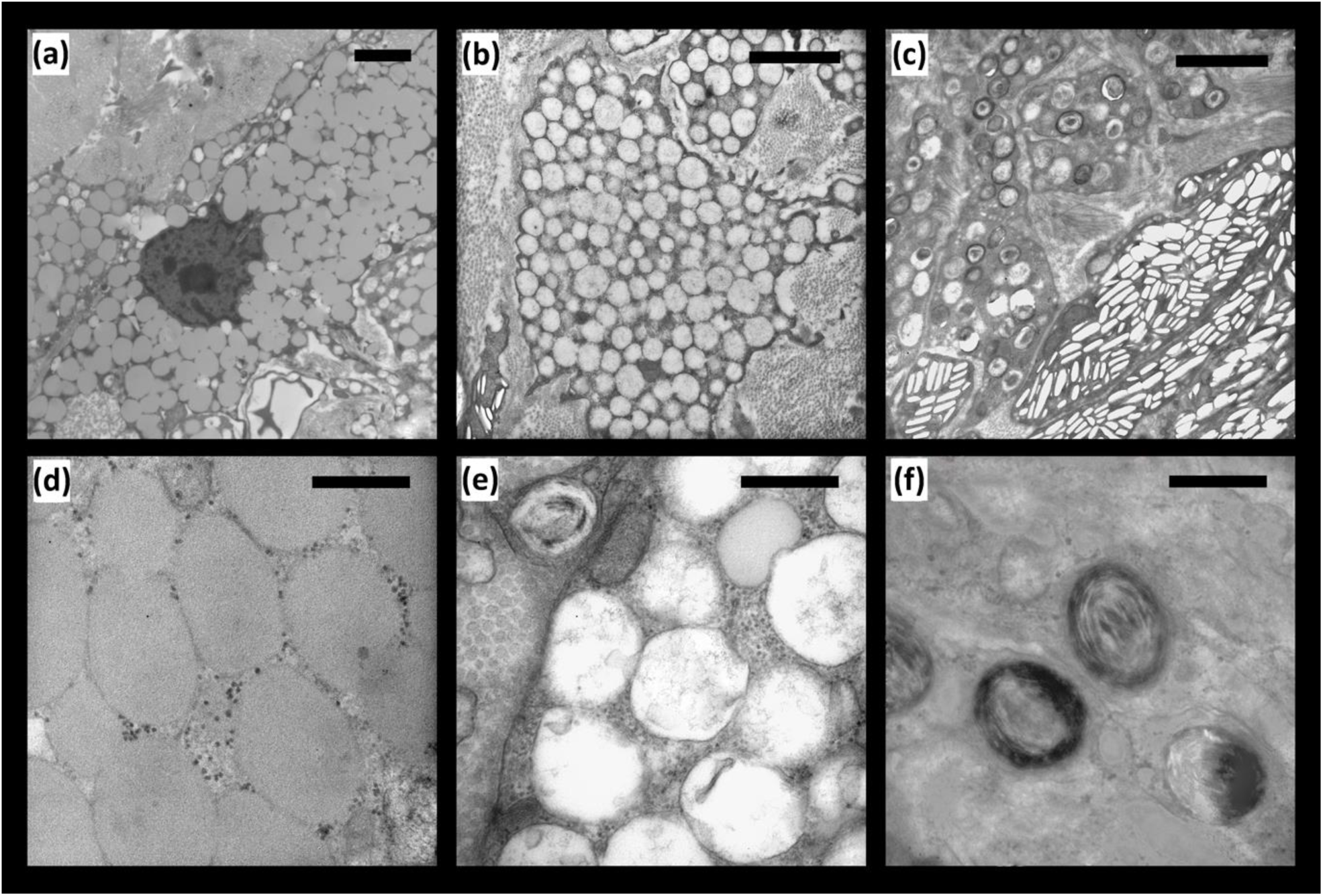
Examples of three different types of vesicles present in the xanthophores of examined turtles. (a) electron dense vesicles present predominantly in the PM of *Trachemys scripta elegans*; (b) vesicles with lower electron density present predominantly in the YP of *Trachemys scripta scripta*; (c) pterinosomes; (d) detail of electron dense vesicles from a), these vesicles lack any internal structure and possibly contain ketocarotenoids; (e) detail of vesicles with lower electron density shown in (b), there is no pronounced structure visible inside these vesicles. In context of chemical analysis, it seems these vesicles contain predominantly yellow carotenoids; (f) detail of pterinosomes shown in (c), typical by concentric lamellae. Scalebar 2µm (a, b, c) and 500nm (d, e, f). Note that magnification of (a) differ.

*P. concinna* has the same pigment cell types in the ventral and dorsal sides of the head. In contrast, the dorsal and ventral regions of the head in the two subspecies of *T. scripta* differ in the distribution of colour-producing pigment cells (Fig. 10). The ventral CBC (Fig. 7 a) and FLBS (Fig. 7 c, d) of *P. concinna* and *T. scripta* contain iridophores together with xanthophores in the dermis. This organization is found also in the PM of *P. concinna* (Fig. 8 e, f; Fig. 7 b). However, the dorsal head regions of both subspecies of *T. scripta*, i. e. yellow YP of *T. s. scripta* (Fig. 8 d), the dark PM of *T. s. scripta* (Fig. 8 c), and the red PM of *T. s. elegans* (Fig. 8 a) contain only xanthophores in the dermis. Another difference among the studied species is that, unlike *T. scripta*, the pigment cells of *P. concinna* do not form continuous layers.

**Figure 10:**
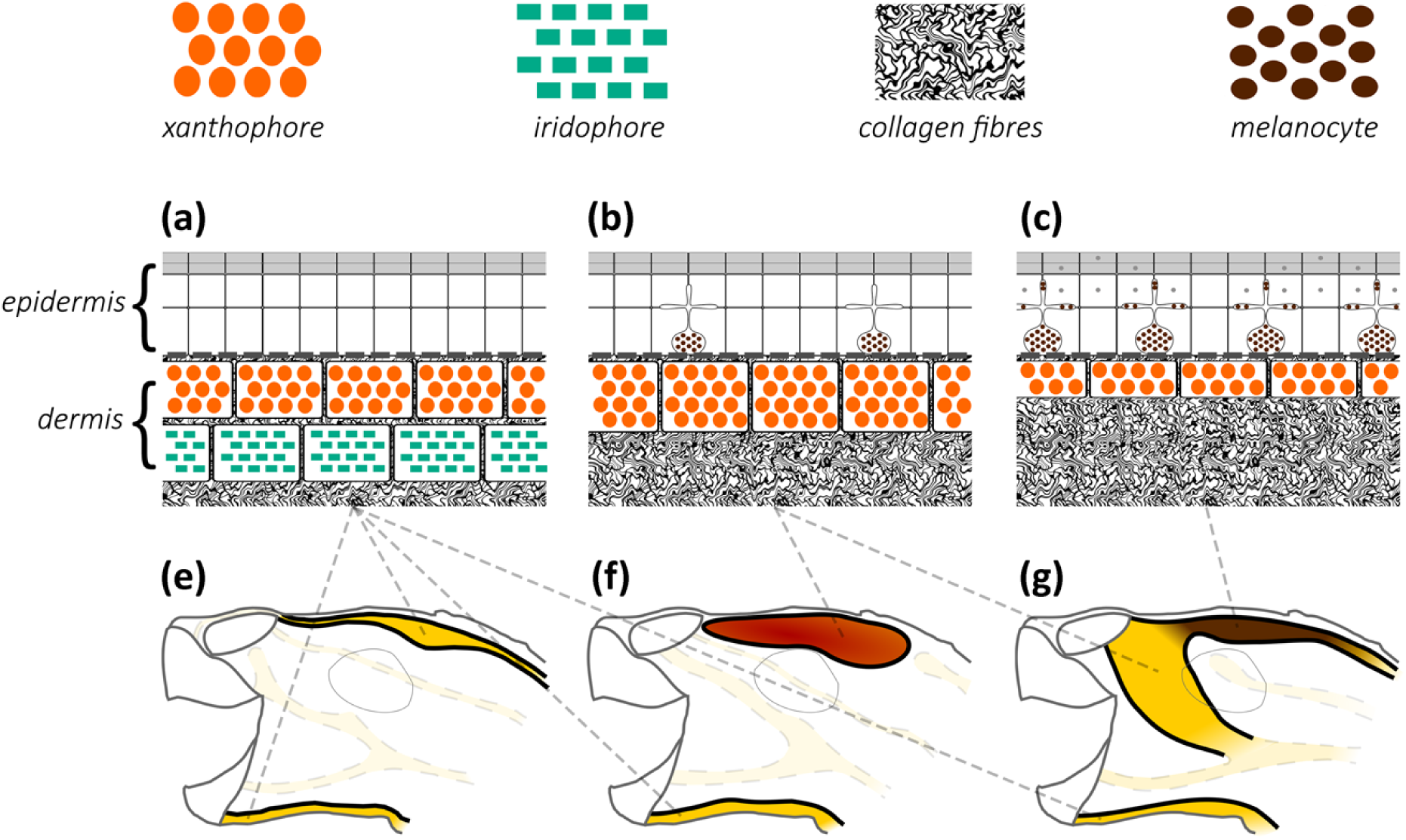
Schematic diagram of cellular composition of colourful region of Emydinae. Ventral chin stripe (CBC), and dorsal postorbital marking (PM) and zygomatic patch (YP) of examined turtles compose of epidermal melanocytes and dermal xanthophores and iridophores. Both ventral and dorsal regions of *Pseudemys concinna* (e) consist of xanthophores and iridophores (a). However, integument of *Trachemys scripta elegans* (f) and *T. s. scripta* (g) contain xanthophores together with iridophores on ventral side. On dorsal side there are only xanthophores in the dermis (b, c) present in both subspecies of *T. scripta*. There are no epidermal melanocytes in neither PM or CBC of *P. concinna* (a), but there are epidermal melanocytes in dorsal integument of *T. scripta*. These epidermal melanocytes are small, without apparent melanosome transferring activity (b) in red PM of *T. s. elegans* and yellow zygomatic region (YP) of *T. s. scripta*. In dark PM region of *T. s. scripta* (c) epidermal melanosomes are enlarged and epidermal keratinocytes contain transferred melanosomes (grey dots). The dermis contains abundant collagen fibres.

Pigment cells form continuous layers whose composition differs between colour regions (Fig. 7, Fig. 8). Bright colour regions contain both single cell type layers (xanthophores) and two cell type layers (xanthophores + iridophores) (Fig. 10). The dark stripes and the background coloration are characterized by the presence of all three cell types but with a predominance of dermal melanocytes. The dermal pigment cells of all studied turtles are embedded in abundant collagen fibers, which in *P. concinna* make up most of the mass of dermis adjacent to the epidermis (compare Fig 4. a, b and Fig 5. a, b).

### 3.3 Reflecting platelets of iridophores

The reflecting platelets present in the iridophores of all regions examined are roughly rectangular (Fig. 11). Unfortunately, measurements of iridophores from the FLBS of *P. concinna* are not included in the results because the platelets were not well preserved. Characteristics and counts of intact reflecting platelets as well as empty holes are summarized in Table 2. Only in the FLBS of *T. s. elegans* and the CBC of *P. concinna* reflecting platelets are mostly parallel to the skin surface (predominant angles of orientation are −1.98° and 2.49°, respectively), while the major axes of reflecting platelets in the CBC of *T. s. elegans* and in the PM of *P. concinna* have average angles differing from zero (30.86° and 68.3°, respectively). Moreover, the reflecting platelets of the FLBS of *T. s. elegans* are more aligned with the skin surface and regularly organized (A/y_0_ = 607.28, FWHM = 59.19°) than other regions examined (CBC of *P. concinna* - A/y_0_ = 10.85, FWHM = 95.3°; CBC of *T. s. elegans* - A/y_0_ = 6.31, FWHM = 106.24°; PM of *P. concinna* - A/y_0_ = 4.91, FWHM = 112.42°) (see supplementary Fig. S3 a, b, c, d for distributions of angles of reflecting platelets for different regions; see measurements of reflecting platelets at [Reference to be added]).

**Figure 11:**
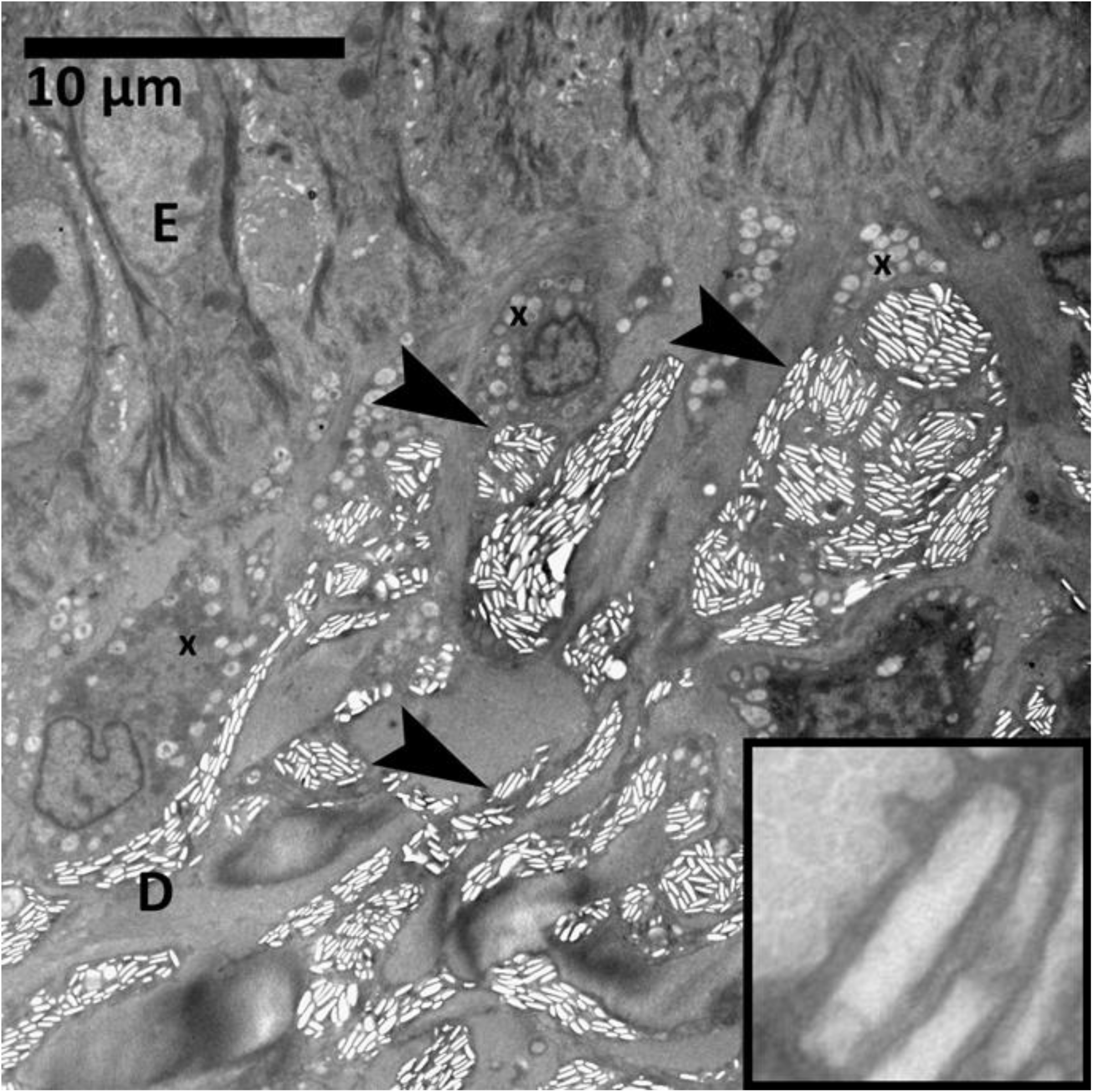
Electromicrograph of iridophores in the turtle integument. E: epidermis; D: dermis; x: xanthophore; arrowheads: iridophore. In the inset is detailed view of reflecting platelet.

Only the FLBS of *T. s. elegans* meets the conditions of the thin layer interference model. Figure 12 a, b shows the predicted reflectivity of iridophores of the FLBS of *T. s. elegans*. The peak of the highest density at the predicted reflected wavelength differs slightly depending on whether intact reflecting platelets or empty holes are measured (see supplementary Fig. S3 e, f, g, h for distributions of thickness of reflecting platelets for different regions based of the two methods of measurement), being 618 nm based on measurement of intact platelets and 637 nm based on measurement of empty holes.

**Figure 12:**
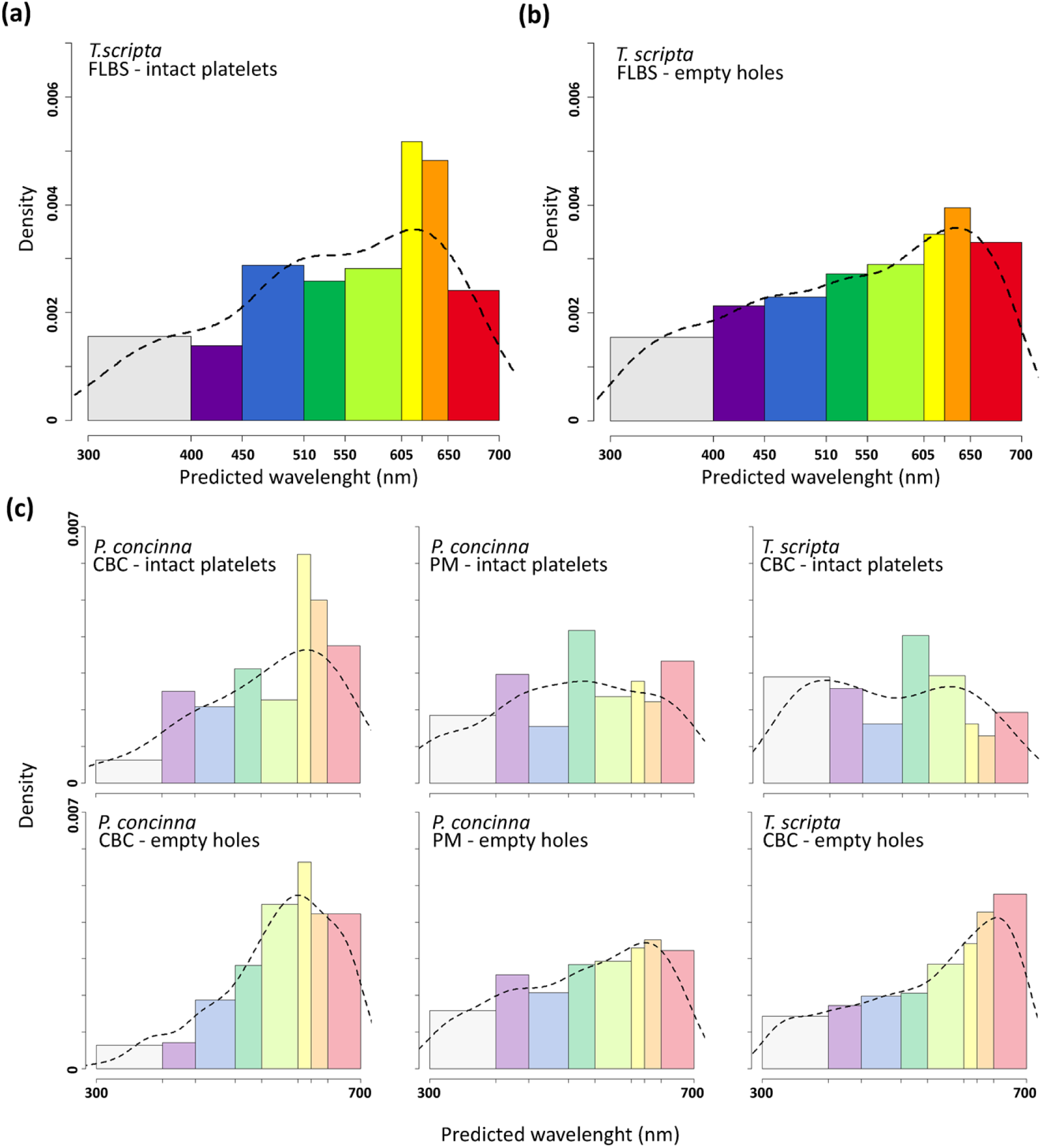
Distribution of predicted reflectivity of reflecting platelets of iridophores. (a) based on size of intact reflecting platelets of FLBS - yellow front limb region of *Trachemys scripta*. (b) based on size of empty holes of FLBS region of T. scripta left after staining of EM sample. Dashed line in (a) and (b) represent shape of reflectance spectra predicted from reflecting platelets properties based on thin layer interference model. (c) distribution of predicted reflectivity of reflecting platelets of iridophores in regions where assumption of thin layer interference model does not seem to be appropriate.

Due to the organization of their platelets, the iridophores in the CBC of *P. concinna* probably function as broadband reflectors. The same probably applies to the iridophores in the CBC of *T. s. elegans* and the PM of *P. concinna*, which in addition to disorganized reflecting platelets also possess platelets of variable size (Fig. 12 c). Only the iridophores of the FLBS of *T. s. elegans* seem capable of intense narrowband colour production.

### 3.4 Fourier analysis of spatial distribution of dermal collagen arrays

The 2-D Fourier power spectra show that none of the single collagen fibre arrays examined (see examples of electromicrographs analyzed in Fig. 13; see output files of Fourier analyses for each elctromicrograph analysed at [Reference to be added]) are organized randomly with respect to wavelengths of light, i. e. the distances between collagen fibres are of the same order of magnitude as wavelengths of light [60]. However, the collagen arrays of *T. s. elegans* show variation in the spatial frequencies of the 2-D Fourier power spectra within regions (supplementary Fig. S4 a, b, c). In contrast, the power spectra of collagen arrays of *P. concinna* show reduced variation in organization within regions (supplementary Fig. S4 d, e, f). Collagen arrays of the PM of *P. concinna* are unambiguously organized in a way which results in small disc pattern of 2-D Fourier power spectra at low spatial frequencies across all micrographs analysed. Collagen arrays of the FLBS of *P. concinna* show two-sided symmetric pattern of 2-D Fourier power spectra at intermediate spatial frequencies rather than a disc pattern, suggesting predominant periodicity in one direction perpendicular to the fibres. In the CBC of *P. concinna* power spectra show a ring pattern at intermediate spatial frequencies, pointing to equivalent nanostructure in all directions perpendicular to the fibres.

**Figure 13:**
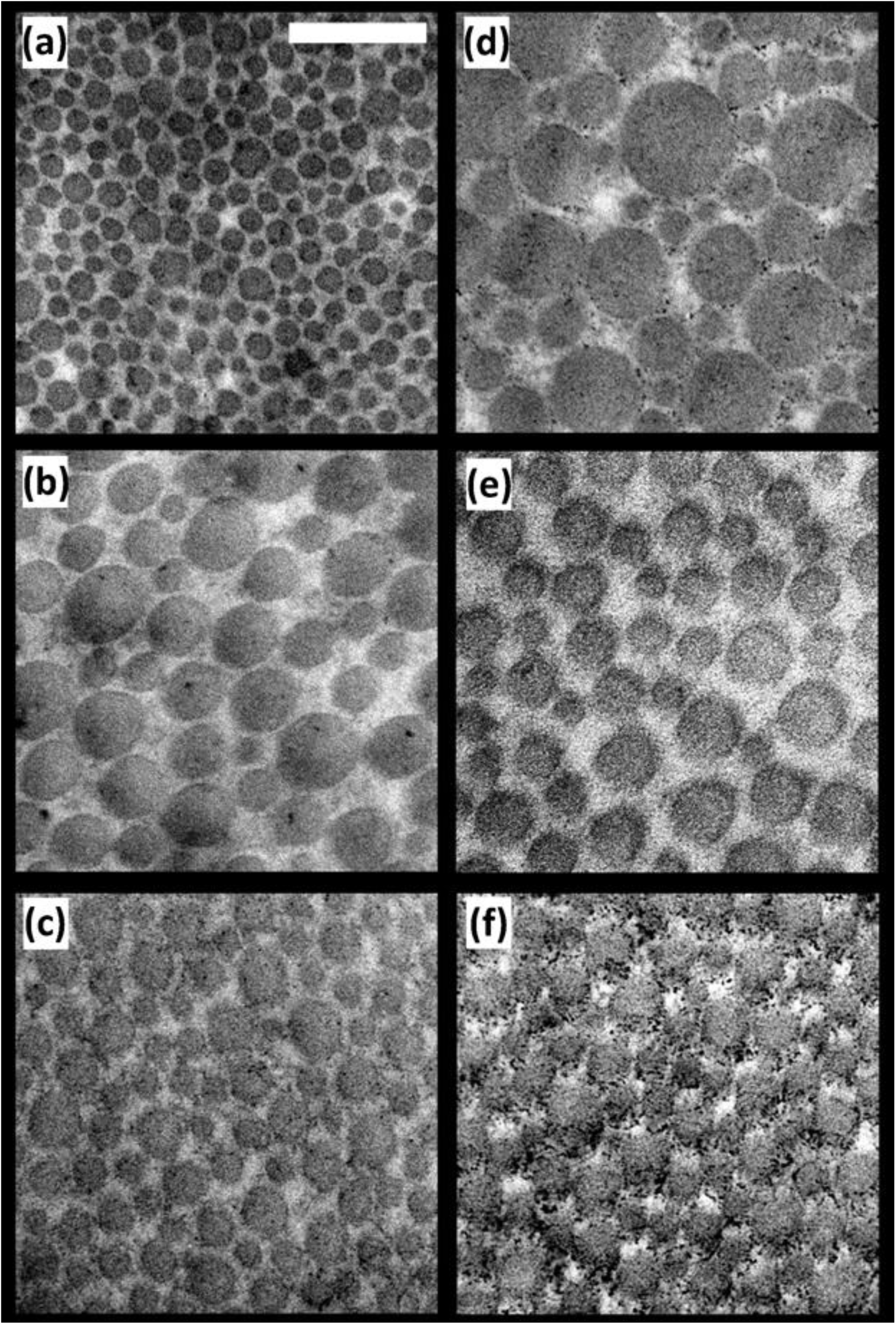
Electromicrographs of details of collagen fibre arrays. (a) Postorbital marking (PM) of the *Trachemys scripta*. (b) Main median chin yellow stripe (CBC) of *T. scripta*. (c) Main bright stripe forelimb stripe of *T. scripta*. (d) PM region of *Pseudemys concinna*. (e) CBC region of *P. concinna*. (f) FLBS region of *P. concinna*. Scalebar represent 200 nm.

Normalized average radial means of the Fourier power spectra for each region of *P. concinna* and *T. s. elegans* are shown in Fig 14. Radial means of the Fourier power spectra of all regions of *T. s. elegans* and the YP of *T. s. scripta* show increased power values in the lowest spatial frequencies (< 0.0034 nm^-1^) compared to spatial frequencies relevant to coherent scattering in the visible light (from 0.0034 to 0.0092 nm^-1^) (Fig. 14 a, b, c; supplementary Fig. S5 a). Radial means of the Fourier power spectra of all regions of *P. concinna* also show increased power values at the lowest spatial frequencies (Fig. 14 d, e, f). These values of spatial frequencies indicate that the largest portion of light reflected by collagen fibres arrays in these regions belongs to the infra-red part of the light spectrum. However, for the CBC of *P. concinna* this peak of low spatial frequencies lies immediately next to the segment of spatial frequencies relevant to coherent scattering in visible light (Fig. 14 e). Radial means of spatial frequencies of collagen fibres of CBC of *P. concinna* relevant to coherent scattering in visible light are in comparison with radial means of all regions of *T. scripta* and the PM of *P. concinna* (Fig. 14 d) relatively elevated. In the FLBS of *P. concinna* collagen fibres are organized at appropriate spatial frequencies (between 0.0034 and 0.0092 nm^-1^) to produce coherent scattering in the visible spectrum (Fig. 14 f).

**Figure 14:**
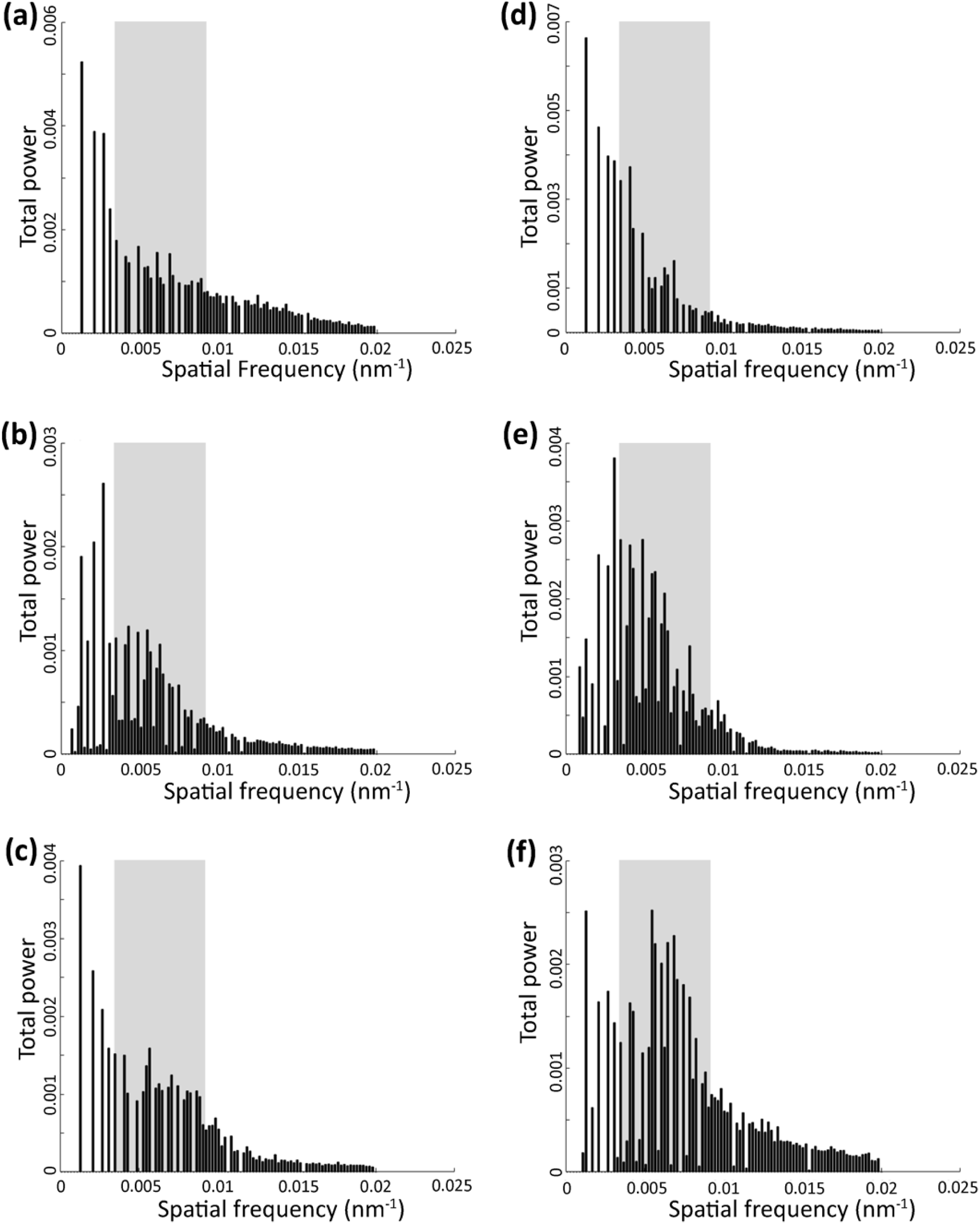
Radial means of the Fourier power spectra of electromicrographs of collagen fibre arrays. (a) Postorbital marking (PM) of the *Trachemys scripta*. (b) Main median chin yellow stripe (CBC) of *T. scripta*. (c) Main bright stripe forelimb stripe of *T. scripta*. (d) PM region of *Pseudemys concinna*. (e) CBC region of *P. concinna*. (f) FLBS region of P. concinna. Grey rectangle denotes spatial frequencies relevant to coherent scattering in visible light, on the other hand lower frequencies (to the left from the grey rectangle) are relevant to scattering in the infra-red part of light spectrum.

Predicted normalized reflectivity of collagen fibre arrays of *P. concinna* and *T. s. elegans* between 300 and 700 nm based on multiple radial means of the Fourier power spectra are shown in Fig. 15 together with normalized measured reflectance spectra. All predicted normalized reflectivity curves have three distinct peaks (360–400 nm, 450–500 nm, and 560–570 nm) except for the YP of *T. s. scripta* which has four peaks (370, 450, 520, and 570 nm; supplementary Fig. S5 b). All predicted normalized reflectivity curves increase beyond 600 nm, but in the CBC of both *T. s. elegans* (Fig. 15 b) and *P. concinna* (Fig. 15 e) there is an additional peak/plateau at 660–680 nm. The overall shapes of the predicted normalized reflectivity of collagen fibre arrays are more similar among regions than among species. The PMs of both species (Fig. 15 a, d) have few distinct predicted peaks with a pronounced peak at 560 nm in the PM of *P. concinna*. The CBC of both species (Fig. 15 b, e) have four significant predicted peaks with the highest at 490–500 nm. The FLBSs (Fig. 15 c, f) were characterized by predicted major peaks in the UV spectrum at 360–380 nm.

**Figure 15:**
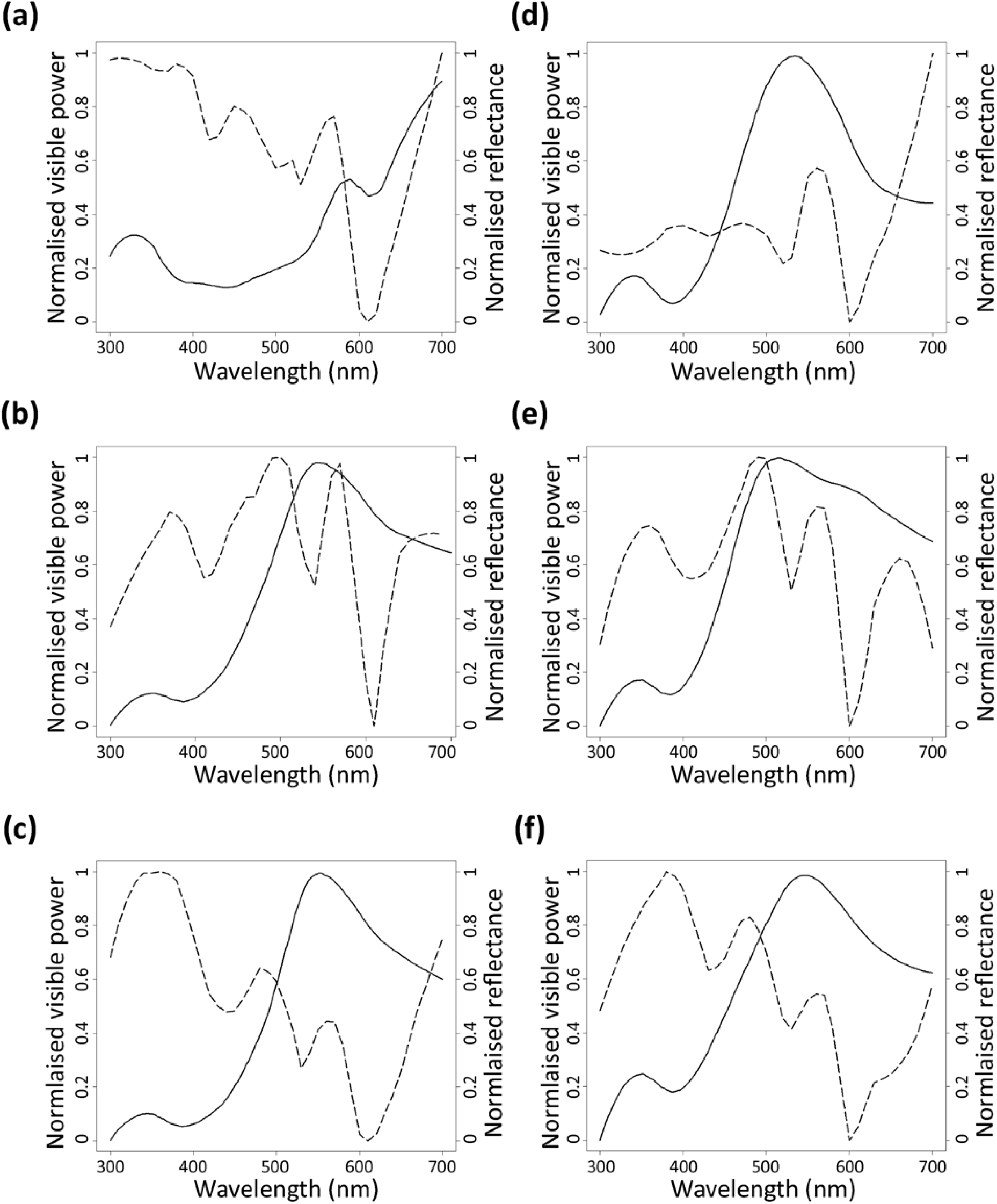
Measured reflectance spectra of the skin regions (solid line) and reflectance spectra predicted by Fourier analyses of dermal collagen fibres (dashed line). (a) Postorbital marking (PM) of the *Trachemys scripta*. (b) Main median chin yellow stripe (CBC) of *T. scripta*. (c) Main bright stripe forelimb stripe of *T. scripta*. (d) PM region of *Pseudemys concinna*. (e) CBC region of *P. concinna*. (f) FLBS region of *P. concinna*. Grey rectangle denotes spatial frequencies relevant to coherent scattering in visible light, on the other hand lowever frequencies (to the left from the grey rectangle) are relevant to scattering in the infra-red part of light spectrum. Note that both measured reflectance and Fourier power are both normalised to have minimum zero and maximum one.

Peaks of the predicted normalized reflectivity of collagen fibre arrays correspond roughly to peaks of normalized measured reflectance spectra. However, there are differences in the overall shapes of the predicted and measured curves. Differences between measured and predicted spectra probably arise due to interactions between colour-producing pigment cells and the intercellular matrix of collagen fibre arrays. Even though the radial means of the Fourier power spectra of the FLBS of *P. concinna* suggest appropriate spatial frequencies to produce colour by coherent scattering, the predicted reflectivity curve does not match the measured reflectance spectra. However, at least in the CBC of *P. concinna* the organization of collagen fibres, radial means of the Fourier power spectra and the congruence of predicted reflectivity curve with measured reflectance spectra suggest a role of collagen fibre arrays in colour production by coherent scattering. The majority of the Fourier power falls into range of presumed infrared part of the light spectrum and contribution of collagen fibre arrays to the overall reflectance is only minor in most other regions. This suggests that the main function of collagen fibres is to reflect infrared radiation and only in some instances have been co-opted to function in colour production.

### 3.5 Pigment contents

Results of pigment content analyses for carotenoids and pteridine derivatives are summarized in Table 4. Examples of UPLC chromatograms resulting from carotenoid content analyses can be found in the supplementary materials (Fig. S1 b, g, h, i, j). The red PM of *T. s. elegans* contains a rich mixture of various carotenoids which, according to UV/VIS absorbance spectra, are predominantly ketocarotenoids. The two most pronounced retention peaks were determined as astaxanthin (RT = 11.8 min, λ_max_ = 471nm, [M+H]^+^ = 597.3944) and canthaxanthin (RT = 13.7 min, λ_max_ = 466, [M]^+^ = 565.4046). Other regions (CBC, FLBS of both species, PM of *P. concinna*, and PM and YP of *T. s. scripta*) had no ketocarotenoids, but contained a relatively complex mixture of hydroxylated xanthophylls, of which lutein (RT = 12.9 min, λ_max_ = 444, [M]^•+^ = 568.4254) and zeaxanthin (RT = 13.1 min., λ_max_ = 449, [M]^•+^ = 568.4253) were by far the most abundant.

**Table 3:**
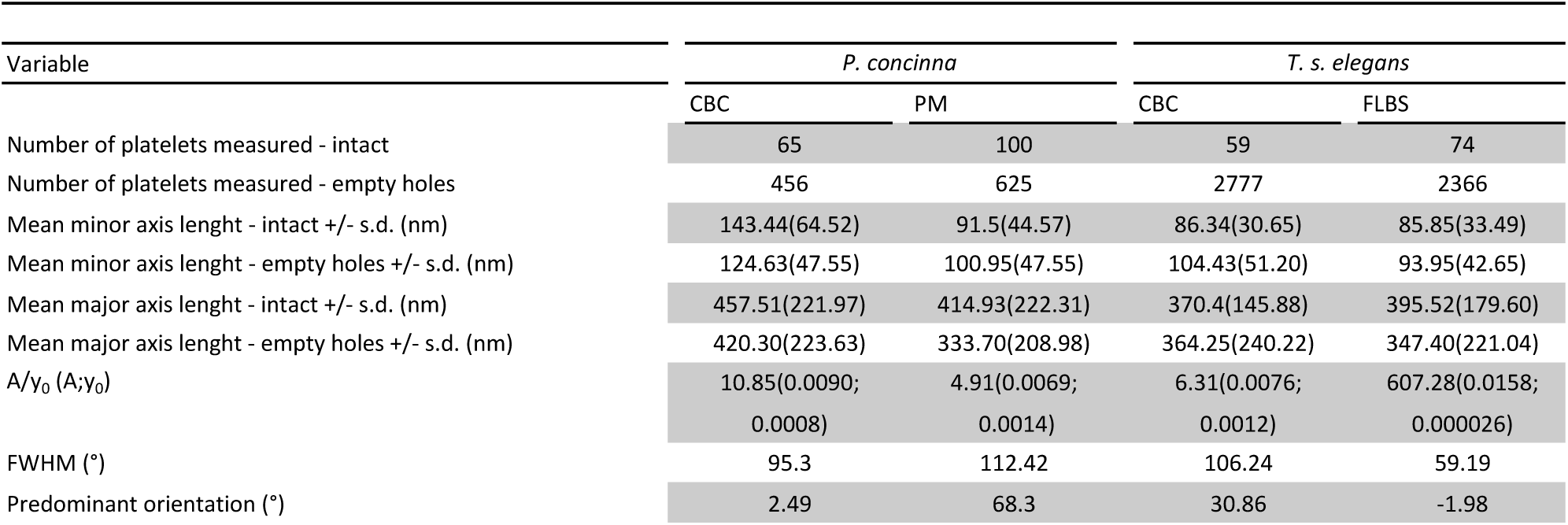
Characteristics of reflective platelets of iridophores in CBC (main median chin yellow stripe), FLBS (main bright stripe of the fore limb) regions of *Pseudemys concinna*, and CBC, PM (postorbital marking) of *Trachemys scripta elegans*. A/y0 ratio determine relative number of reflecting platelets parallel to the skin surface. A: amplitude of Gaussian curve above the background of randomly oriented platelets; y0: background level of randomly oriented platelets; FWHM: full width of half maximum of the peak of Gaussian curve fitted to the data, i. e. width of angular distribution of reflecting platelets.

**Table 4:** Contribution (✔ present/ ✖ absent) of different types of carotenoid and pteridine derivatives in regions of integument in studied taxa (*Pseudemys concinna, Trachemys scripta elegans, Trachemys scripta scripta*). CBC: main median chin yellow stripe; FLBS: main bright stripe of the fore limb; PM: postorbital marking; YP: yellow zygomatic patch.

| Taxon | Region | Pteridine Derivatives |  |  |  |  |  |  |  | Predominant Carotenoids |  |  |  |  |
| --- | --- | --- | --- | --- | --- | --- | --- | --- | --- | --- | --- | --- | --- | --- |
|  |  | 6-biopterin | pterin-6-COOH | Erythropterin | Isoxanthopterin | Leucopterin | L-sepiapterin | D-Neopterin | Pterin | Xanthopterin | Astaxanthin | Canthaxanthin | Lutein | Zeaxanthin |
| *P. concinna* | CBC | ✓ | ✓ | ✗ | ✗ | ✓ | ✗ | ✓ | ✗ | ✗ | ✗ | ✗ | ✓ | ✓ |
|  | FLBS | ✓ | ✓ | ✗ | ✗ | ✓ | ✗ | ✓ | ✗ | ✗ | ✗ | ✗ | ✓ | ✓ |
|  | PM | ✓ | ✓ | ✗ | ✗ | ✓ | ✗ | ✓ | ✗ | ✗ | ✗ | ✗ | ✓ | ✓ |
| *T. s. elegans* | CBC | ✓ | ✓ | ✗ | ✓ | ✓ | ✗ | ✓ | ✗ | ✓ | ✗ | ✗ | ✓ | ✓ |
|  | FLBS | ✓ | ✓ | ✗ | ✓ | ✓ | ✗ | ✓ | ✗ | ✓ | ✗ | ✗ | ✓ | ✓ |
|  | PM | ✓ | ✓ | ✗ | ✓ | ✓ | ✗ | ✓ | ✓ | ✓ | ✓ | ✓ | ✗ | ✗ |
| *T. s. scripta* | CBC | ✓ | ✓ | ✗ | ✓ | ✓ | ✗ | ✓ | ✗ | ✓ | ✗ | ✗ | ✓ | ✓ |
|  | FLBS | ✓ | ✓ | ✗ | ✓ | ✓ | ✗ | ✓ | ✗ | ✓ | ✗ | ✗ | ✓ | ✓ |
|  | YP | ✓ | ✓ | ✗ | ✓ | ✓ | ✗ | ✓ | ✗ | ✓ | ✗ | ✗ | ✓ | ✓ |

Examples of SRM chromatograms of pteridine analyses are given in the supplementary material (Fig. S6). Out of nine pteridine derivatives tested, seven were found in the integument of the turtles analysed. L-sepiapterin and erythropterin were not detected in any sample. All other pteridine derivatives, i. e. 6-biopterin, pterin-6-COOH, isoxanthopterin, leucopterin, D-neopterin, pterin, and xanthopterin, were found together only in the PM of *T. s. elegans*. The yellow CBC and FLBS of both lineages of *T. scripta* and the YP of *T. s. scripta* contained all types of pteridine derivatives except pterin. Only four types of pteridine derivatives, 6-biopterin, pterin-6-COOH, leucopterin, and D-neopterin, were detected in the CBC, FLBS, and PM of *P. concinna*.

The red PM of *T. s. elegans* is unique among the examined region because it contains ketocarotenoid and seven pteridine derivatives, including pterin, which was found in no other region. The ventral (CBC, FLBS) and dorsal yellow regions (YP) do not differ in pigment contents in *T. scripta*. All yellow regions of *T. scripta* (CBC, FLBS, YP) and *P. concinna* (CBC, FLBS) have the same carotenoid contents. On the other hand, pteridine derivatives are more diverse in the yellow regions of *T. scripta* than in those of *P. concinna*. Two additional pteridine derivatives found in the integument of *T. scripta* (isoxanthopterin and xanthopterin) are known to produce yellow colour. The integument of *P. concinna* contains the same pigment types in every region examined (PM, CBC, FLBS).

## 4. Discussion

Although they are not known for their vivid, conspicuous body coloration, many turtles are in fact as colourful as other vertebrates [69]. However very few studies have addressed the mechanisms of turtle colour production [27,29,30]. In fact, there is more information on colour production in some fossil extinct taxa than in extant turtles [70–76]. Research on the functional role of turtle chromatic signals is also limited [41], although it is known that at least some species change colour during the reproductive season [19]. Given the revised phylogenetic position of turtles as part of archelosauria [22,77–79], studying turtle coloration and colour-producing mechanisms crucial to understanding the evolution of colour and colour-producing mechanisms in vertebrates in general.

In this study we analysed the pigment cell organization of the integument, the ultrastructure of colour-producing elements, and the chemical nature of pigments that produce skin colour in two freshwater turtles, *Pseudemys concinna* and *Trachemys scripta*, with contrasting courtship behaviour. We found striking interspecific differences in the organization of the chromatophores found in the dorsal and ventral head surfaces. *Trachemys scripta* shows a different chromatophore composition in the ventral and dorsal sides of the head, whereas *P. concinna* shows similar chromatophore composition on both sides. Also, there are more pigment types present in the colour patches of *T. scripta* than in those of *P. concinna*.

Turtles employ colour producing mechanisms well known in amphibians and lepidosaurs, but not in other extant archelosaurs (crocodiles and birds). In addition to xanthophores and melanophores previously reported in turtles [27], and crocodiles [80], we found abundant iridophores containing rectangular reflecting platelets in yellow skin of both species. To our knowledge, this is the first report of iridophores playing a role in integumental colour production in any archelosaurian species [80,81]. Carotenoids are involved in colour production in the feathers of birds [82], while pteridines are responsible for the colour of the iris of some birds [83] and perhaps also the feathers of penguins [84]. However, our pigment analyses suggest that both pteridine derivatives and carotenoids are involved in the production of the yellow-red skin colours of turtles, which had not been clearly documented in any archelosaur to this date.

We also show that abundant collagen fibre arrays in the skin may serve to prevent infra-red light from penetrating into the body of turtles, but in some cases, they may be capable of reflecting visible wavelengths of light as well. Taken together, these results suggest that turtle coloration and colour-producing mechanisms are more complex than previously thought.

### 4.1 The coloration of deirocheline turtles

Our results show that the conspicuous yellow stripes in the head and limbs of deirocheline turtles exhibit, in addition to a primary reflectance peak responsible for the human-perceived yellow hue, a small secondary peak of reflectance in the UV range of the spectrum. This confirms the results of several previous studies [30,85]. However, we did not find support for the spectral shapes reported by Wang et al. [86]; in fact, we contend that the 372 nm peak that these authors found in all their reflectance spectra (see their Figure 1 [86]) is an artefact possibly caused by the illumination source. As turtles are capable of seeing into the UV range [87,88], the small peak of UV reflectance may account for chromatic variation among different individuals and contribute to the conspicuousness of the yellow stripes, at least during close-range interactions with conspecifics [30]. It is also possible that the pattern of alternating dark and light stripes may provide camouflage through background matching (e.g. when viewed against underwater vegetation) or as disruptive coloration creating false edges and visually breaking the outline of the body, thus hindering detection of turtles by predators [89].

Our results reveal interspecific differences in colour-producing mechanisms at multiple levels: differences can be found in the distribution of chromatophores along the dorsoventral body axis, in the ultrastructure of pigment-bearing vesicles of xanthophores, and in the chemical nature of pigments involved in colour production. As for the distribution of chromatophores, *Trachemys scripta* has both xanthophores and iridophores in the yellow-red regions of the ventral side of the head, but only xanthophores on the dorsal side. Such dorso-ventral patterning is lacking in *P. concinna* which has xanthophores and iridophores on both sides of the head (Fig. 10). As iridophores increase the overall reflectance of a patch of skin [3,20,90], their presence on the ventral side of the head may provide some sort of countershading to enhance crypsis ([91–93]), or may be related to signalling (see below). The fact that both species of turtles also have a light plastron, but a dark coloured carapace is consistent with the hypothesis that the increased reflectance caused by the presence of iridophores on the ventral side of the head might serve a camouflage function. The background head coloration (DHC), which has been previously reported to change in response to substrate colour [36,94–96], is produced in both *T. scripta* and *P. concinna* by epidermal melanosomes and a combination of all three dermal pigment cell types (Fig. 7 e, f). However, we found epidermal melanocytes in the PM of both lineages of *T. scripta* (Fig. 5, Fig. 8 e, f), but not in the PM of *P. concinna* (Fig. 4, Fig. 8 a, b, c) or in other yellow regions of the examined turtles (Fig. 7 b, d).

The red and yellow regions of turtle skin contain at least three morphologically distinct types of vesicles in xanthophores. Two of these correspond to what have been previously [68,97,98] identified as carotenoid vesicles, and differ in electron density, shape and size (Fig. 9 a, b, d, e). The third type of xanthophore vesicles are pterinosomes, likely containing pteridine derivatives (Fig 9 c, f). These are recognizable by their prominent concentric lamellae [67,68]. Xanthophores containing oval carotenoid vesicles with high electron density (Fig. 9 a, d) are abundant only in the red PM of *T. scripta elegans*, while xanthophores containing round carotenoid vesicles with lower electron density (Fig. 9 b, e) and/or pterinosomes (Fig. 9 c, f) are found in yellow regions of both species. We assume that differences in the ultrastructure of carotenoid vesicles reflect differences in their pigment content, as in the case of red and yellow retinal oil droplets [99] known to differ in carotenoid content [31,100,101]. However, similar structures have been previously described as vesicles in different stages of maturation [102]. Further study using appropriate methods is necessary to unambiguously determine the types of pigments contained in vesicles of different ultrastructure.

Continuous layers of iridophores are found only in *T. scripta*. Moreover, the reflecting platelets of iridophores from the CBC of *T. scripta* are highly organized with predicted reflected wavelengths between 618 and 637 nm (Fig. 12 a, b). Other regions containing iridophores, i. e. yellow FLBS of *T. scripta* and all yellow regions of *P. concinna*, do not have organized platelets and therefore function as broadband reflectors which should enhance the overall brightness/luminance of the corresponding skin patch. Previous reports have described iridophores with randomly organized platelets in red and white skin of *Phelsuma grandis* where they produce broadband reflection [16]. In contrast, reflecting platelets in orange and yellow skin regions of *Uta stansburiana* [57], *Sceloporus undulatus*, and *S. magister* [56] are highly organized to reflect orange and yellow wavelengths. Even though broadband reflecting iridophores do not influence perceived colour, they may increase the brightness of colour regions and their overall conspicuousness to observers [3,103].

Unlike all yellow regions of both species, which contain xanthophylls lutein and zeaxanthin, the red PM of *T. s. elegans* contains the ketocarotenoids astaxanthin and canthaxanthin (Table 4, Fig. S1). Such difference is in partial agreement with previous analyses of carotenoid contents in yellow and red regions of *T. s. elegans* and *Chrysemys picta* [30]. Two major classes of carotenoids have been described in the integument of these turtles: short wavelength absorbing apocarotenoids and longer wavelength absorbing ketocarotenoids [30]. In the yellow chin of *C. picta* only apocarotenoids are present, whereas the orange neck and leg contain apocarotenoids and ketocarotenoids. Previous reports indicated that apocarotenoids are abundant in tissue from yellow chin and yellow neck stripes of *T. scripta*, its “orange” postorbital region containing only ketocarotenoids [30]. Our results, however, suggest a major role of xanthophylls, rather than apocarotenoids, in yellow colour production in *T. scripta* and *P. concinna*. Xanthophylls have been shown to produce yellow coloration in various vertebrates [82,104,105]. In *C. picta* increased ingestion of the xanthophyll carotenoid lutein results in an increase in the yellow and red chroma of yellow and red skin patches [28]. In contrast, it is unclear whether apocarotenoids participate in colour production in animals. Apocarotenoids are cleavage products of carotenoids [106–108]. Transparent and clear cone oil droplets in the retina of birds contain the apocarotenoid galloxanthin, but yellow oil droplets contain the xanthophyll zeaxanthin and red oil droplets contain the ketocarotenoid astaxanthin [109]. The unique yellow colour of the *macula lutea* in primates is due to the accumulation of xanthophylls in the cells of the retina via down-regulation of their cleavage pathway to apocarotenoids [110]. Various carotenoid precursors give rise to a large diversity of apocarotenoids [106], but specific types of apocarotenoids have not been reported in previous studies of turtle coloration [30]. The role of apocarotenoids in turtle coloration will thus remain unclear until the relationship between precursor carotenoids in the skin and apocarotenoids is elucidated.

Steffen et al. [30] suggested that the yellow-red skin of *C. picta* and the yellow skin (but not the red skin) of *T. scripta* contain small amounts of pteridine derivatives. Our results show that the opposite is true (Table 4). The red PM shows more types of pteridine derivatives than the yellow skin in *T. scripta*. Yellow skin in *P. concinna* contains only a limited number of pteridine derivatives in comparison with all regions of *T. scripta*. Our results thus show that the red PM of *T.s. elegans* is more elaborate in terms of its pteridine derivative contents than all the yellow regions studied herein which contrasts with the previous study of Steffen et al. [30]. The yellow and red colours of the turtles examined here result from an interplay between carotenoids and pteridine derivatives.

Whether the differences in colour and colour-producing mechanisms among deirocheline turtles are of functional significance has yet to be determined. However, it is already well established that turtles are capable of distinguishing different yellow and red colours [34,87,88] and it has been suggested that body coloration plays a role in mate choice in emydid turtles [30,34,35]. Furthermore, there is some evidence that the red and yellow regions of *T. scripta* could play a role in signalling individual quality based on the correlation of colour and induced immune response [111]. Moreover, the spectral properties of colour patches of *C. picta* is influenced by the amount of ingested carotenoids [28] which is one of the basic assumption of individual quality signalling [112–114].

Is there a relationship between chromatophore distribution and male-female position during courtship? *Trachemys scripta* and *P. concinna* differ in the distribution of chromatophore and pigment types along the dorsoventral body axis (Fig. 10). The presence of xanthophores together with iridophores in the ventral CBC, but not in the dorsal PM, as well as the presence of epidermal melanocytes in the PM, may enhance the cryptic appearance of *T. scripta* in aquatic environments (see above). It is not clear, however, why *P. concinna* does not possess a similar distribution of pigment cell types. Instead, *P. concinna* has no epidermal melanosomes in the PM, and xanthophores and iridophores are distributed dorsally (PM) and ventrally (CBC). This should make *P. concinna* rather conspicuous when viewed from above. Many animals solve the trade-off between making themselves conspicuous to conspecifics and avoiding detection by predators by restricting bright colours to body surfaces that are exposed to conspecifics but remain hidden to predators (called signal partitioning; e.g. [115,116]). This is often the case with sexual signals which are made visible to mates during courtship [38,39]. Female mate choice is presumably widespread among emydid turtles [117]. Males of *T. scripta* court females in a face-to-face position, so that both the dorsal and ventral sides of the head as well as the limbs of the male are simultaneously exposed to the female. On the other hand, males of *P. concinna* swim above the female during the final stages of courtship [44]. In this position it may be more difficult for the female to perceive simultaneously the dorsal and ventral sides of the male’s head. Thus, the location of the potentially brighter surfaces on the head of the males does not match predictions based on the relative positions of males and females during courtship, at least in the two species examined here.

The red and yellow body regions of *Trachemys scripta* contain a more diverse palette of pteridine derivatives than the yellow regions of *P. concinna* (Table 4). This evidence suggests another possible link between coloration and courtship position. The face-to-face position may facilitate female assessment of male coloration in *T. scripta*, where red and yellow colours could function as signals conveying information about individual male quality. Conversely, the relatively simple pigment contents of species that use the swim-above position would suggest a reduced role for male head coloration in female mate assessment. Thus, the limited ability of male to simultaneously contrast different body regions against female’s visual apparatus may constrain the differential elaboration of various body regions in *P. concinna*.

Female mating preference for colourful males often results in sexual dichromatism, with brightly coloured males and comparatively dull females [118]. Males of *T. scripta* have brighter CBC and FLBS than females, whereas in *P. concinna* sexual differences are less pronounced and restricted to the CBC (Fig. 2, see also supplementary materials and Table S4). This contrasts with a previous study reporting no intersexual differences in coloration in *T. scripta elegans* [84]. Other genera of deirocheline turtles court in a face-to-face position, specifically those in the genera *Chrysemys, Graptemys*, and *Malaclemmys*. It seems likely that other factors besides courtship position contribute to interspecific variation in colour and colour-producing mechanisms in deirocheline turtles, including habitat type, predation and evolutionary history. As mating strategies are relatively well studied in this group, it may be relatively straightforward to test hypothesis about the role of natural and sexual selection in shaping the evolution of colour across the Deirochelinae. Finally, it is important to note that not only function but also differences in proximate developmental processes may be responsible for differences in coloration between *T. scripta* and *P. concinna* which may not necessarily relate to visual signalling or crypsis, but rather to the intrinsic self-organizing properties of pigment cells [102,119–122] Further studies are needed to clarify the development of pigment pattern formation in turtles [123,124].

### 4.2 The function of the reflective shield of collagen fibres

Results of Fourier analyses show that turtle collagen fibres reflect in the infrared part of the spectrum in all regions examined (Fig. 14). Long wavelengths of light that pass through the xanthophores and iridophores may be absorbed by the melanophores, or they may be reflected back to the surface by the underlying collagenous connective tissue when melanophores are absent or scarce [125]. In many fish and amphibian larvae, the collagen fibres form subepidermal collagenous lamella overlying the dermal chromatophores [20,126,127]. In lizards and snakes collagenous connective tissue is found in the *superficial fascia* between the integument and the underlying muscle [125], but in other animals such as frogs [128], mammals [17], or birds [18] collagen fibres are part of the dermis. Here we show highly abundant dermal collagen fibre arrays located below the chromatophore layers in turtles (Figs. 4, 5). Unlike lizards [16,57,97] turtles lack a layer of dermal melanophores in the red-yellow skin regions. Turtles bask to thermoregulate and overheating may represent an immediate risk for them [129], but basking could also have a social function [130] for which it may be advantageous to accumulate heat at a slower rate. Therefore, the function of dermal collagen fibres may be to prevent long infrared wavelengths from reaching the body core to avoid overheating. A thermoprotective function of colour-producing structures has been recently suggested for the deep iridophores found in chameleons [55] or plumage of birds [131]. The role of melanin in thermoregulation is widespread across a broad range of organisms [132]. Thus, it seems that colour-producing elements are commonly acquired to thermoregulate and, conversely, skin structures that play role in thermoregulation may in some instances be co-opted to produce visible colour.

Dermal collagen may appear achromatic [128], but colour production by coherent scattering on nanostructured collagen fibre arrays has been previously reported in mammals [17] and birds [15,18]. In mammals the colour produced by scattering on collagen fibres is limited to the blue part of the light spectrum [17], but in birds other colours, such as orange in *Trogopan coboti*, are produced by coherent scattering on collagen fibre arrays [18]. The yellow FLBS of *P. concinna* shows sufficient nanostructural organization to produce yellow colour by coherent scattering (Fig. 14 f). In most of the examined regions of *T. scripta* and *P. concinna* there is also small peak in the UV part of the spectrum which is predicted by Fourier analyses. Our results predict colour production by collagen fibres in all regions (Fig. 15). However, in most regions the collagen fibers do not conform to the conditions of coherent scattering or the predictions do not match the measured reflectance spectra very well. Even though the reflected light by collagen fibre arrays would be achromatic their presence could still affect the reflectance spectra. It has been shown theoretically that xanthophores have the highest influence and iridophore the lowest influence on perceived colour when the melanophore layer is absent and reflected light off the collagen fibres passes back to the surface [3]. As in the CBC and FLBS of *P. concinna* chromatophores are scarce, it is possible that collagen fibres play a major role in colour production in this species.

### 4.3 Significance for vertebrate skin colour evolution

Most of vertebrates do produce various colours by superposition of different dermal pigment cells, however homeothermic tetrapods, the birds and the mammals, acquire their coloration mainly due to the physiological actions of epidermal melanocytes [133]. The archelosaurian clade, comprising turtles, crocodiles, and birds [22], has undergone dramatic evolutionary changes affecting colour production mechanisms [134]. Some mechanisms of colour production, such as melanogenesis, were modified perhaps with the acquisition of feathers [134], while others, such as the use of ketocarotenoids for red coloration, may have been acquired early during the evolution of the archelosaurian clade [31]. Iridophore and xanthophore differentiation or colour production by pteridine derivatives have been lost or reduced during the evolution of the archelosaurian clade [84,133,135–138]. However, cells with lipid vesicles suspected to contain pigments similar to xanthophores have been reported in both the epidermis and the dermis of non-feathered bare skin of some bird taxa [18]. Crocodiles develop iridophores and xanthophores that participate together with dermal melanophores/melanocytes in the production of brown-yellow background colour, but the iridophores of crocodiles do not poses differentiated reflecting platelets [80]. Previous results suggest that crocodiles may have gone through a nocturnal bottleneck after their divergence from the lineage leading to birds while reducing their colour discrimination capacity [139], and thus may not represent an ideal group to study colour evolution in the archelosaurian clade. Here we show that turtles possess all three types of dermal pigment cells (xanthophores, iridophores, and melanocytes), shared with fish [140,141], amphibians [142,143], and lepidosaurs [144–146]. Analyses of the pigments involved in the production of yellow-red colours in turtles suggest that pteridine derivatives contribute, together with carotenoids, to these colours, which was not previously documented in non-avian archelosaurs, but has been reported in fish [112], amphibians [147], and lizards [104,148]. We show that dermal collagen fibres participate in colour production in turtles which has been reported in mammals [17] and birds [18] and adult amphibians, but not in fish [20,126], amphibian larvae [127], and lizards [125]. Turtles have also been reported to produce colour by hyperproliferation of keratin in epidermal keratinocytes [19]. The keratin matrix is known to play a vital role in the coloration of birds’ feathers [149,150]. The evidence thus suggests that studies of turtle coloration are crucial to understanding the evolution of chromatic diversity in the archelosaurian clade and in tetrapods/vertebrates in general, because turtles seem to employ most of the mechanisms of colour production known in other groups of vertebrates (Fig. 16). Moreover, the field of paleo-colour, which aims to reconstruct the coloration of extinct taxa, has grown dramatically in recent years, providing some of the most remarkable breakthroughs in our understanding of the evolution of animal coloration [72,73,134,151–153] and turtles, given their key position in the phylogenetic tree [77–79] therefore may provide important insights for the proper interpretation of paleontological data.

**Figure 16:**
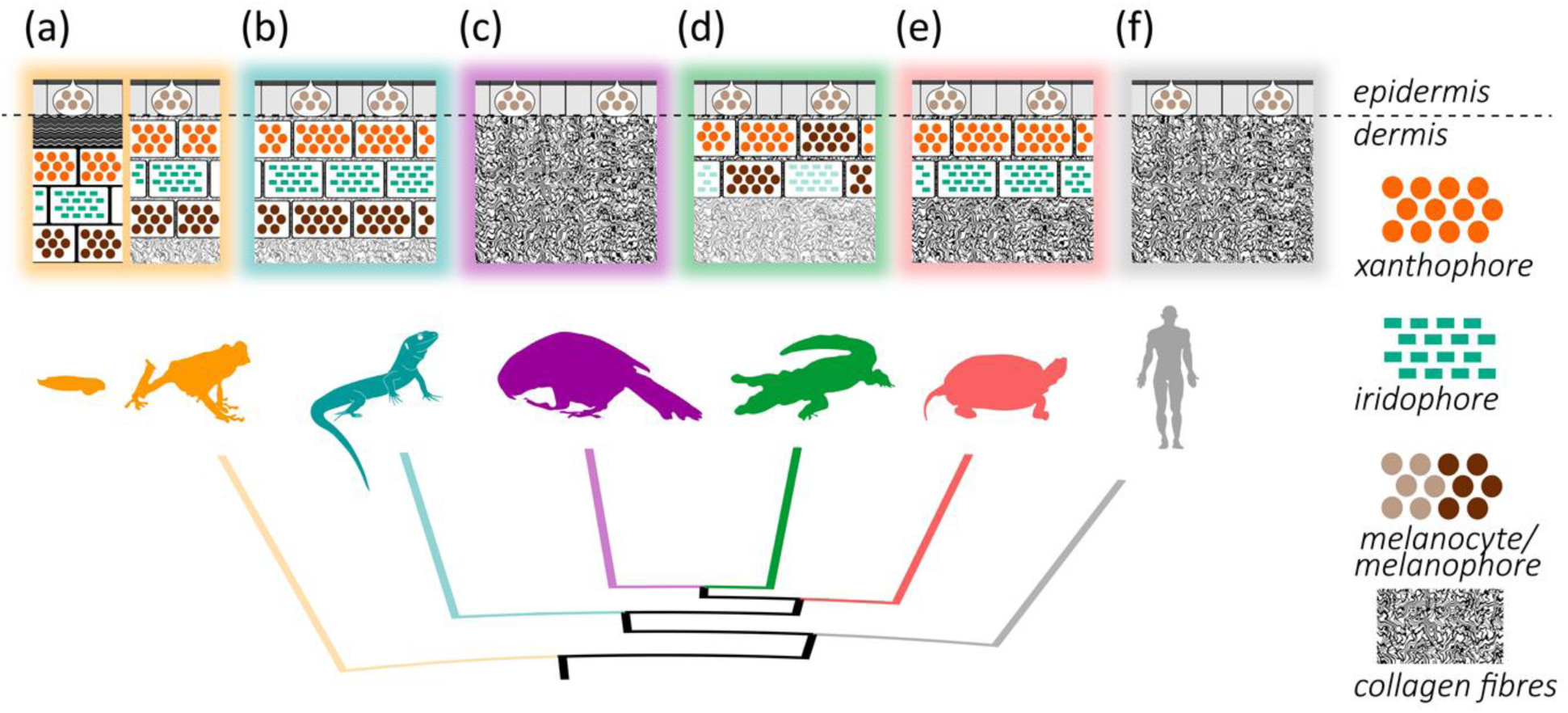
Phylogeny of extant tetrapods [156] illustrating distribution of composition of colour-producing elements in the skin of tetrapods. All of the tetrapods may pose epidermal melanocytes in the epidermis. (a) composition of colour-producing elements of amphibian larvae (on the left) is similar to that of fish. Basal lamina is underlined by layer of subepidermal collagen fibres beneath which are xanthophores, iridophores and melanophores/melanocytes located in the dermis. Adults of amphibians dermal xanthophores, iridophore and melanophores that may form dermal chromatophore units. (b) lepidosaurs pose similar organization of colour-producing elements to that of adults of amphibians. (c) birds do not pose any additional pigment cell type to the epidermal melanocytes. Birds are known to produce colour by organized collagen fibre arrays. Means of colour production of plumage are not depicted. (d) crocodiles pose dermal pigment cells: xanthophores, iridophores, melanocytes, that have not been reported to form continuous layers. Iridophores of crocodiles have been reported to not contain rectangular reflecting platelets. (e) turtles do pose in dermis continuous layers of single xanthophores in some regions, other regions may be characterized by presence of continuous double layer of xanthophores together with iridophore in the dermis, which have not been found underlined by melanophores/melanocytes in neither of the skin regions. Single continuous layer of dermal melanocytes has been found in turtles. Dermal collagen fibres participate in colour production of turtles (f) mammals do not pose any additional pigment cell type to the epidermal melanocytes. Mammals are known to produce colour by organized collagen fibre arrays.

### 4.4 Future directions

More research is needed to further examine if the evolution of coloration in turtles is constrained by the position during courtship and to test the idea of modular evolution of colour ornaments in turtles. Future studies should consider a broader range of species of turtles and apply comparative cophylogenetic methods to map ancestral states of coloration, its mechanisms of production, and relevant functional traits such as courtship behaviour.

In this study we have not addressed the ontogeny of colour-producing mechanisms, as our knowledge of the early development of chromatophores in turtles is still scarce [123]. Also, the postnatal ontogeny of colour in turtles is studied insufficiently and long-standing questions remain unresolved, e. g. melanophores in *T. scripta* are found mostly in the dermis while older “melanistic” males also have elaborate epidermal melanocytes [48], which raises the question how melanophores make the transition from one skin layer to another.

Our results also suggest that early archelosaurs share colour-producing mechanisms with amphibians and lepidosaurs but also used novel ways to produce skin colour such as scattering on dermal collagen fibres. Studies of colour-producing mechanisms on a broad phylogenetic scale are critical to clarify the evolution of colour-producing mechanisms across vertebrates and should be encouraged in the future.

Turtles are a critically endangered group with 61% of species threatened or already extinct [154], which may seriously limit the opportunities to collect biological samples. On the other hand, *T. scripta* is considered one of the most widespread invasive alien species [155] and intensive management measures are taken to control its introduced populations. Pertinent culling of individuals from introduced species offers an excellent opportunity to collect rare biological samples which should not be missed.

## Acknowledgments

We are thankful to Piscífatoria del Palmar for the exceptional hospitality. This study would not be possible without Vicente Sancho and Ignacio Lacomba who pioneered turtle conservation in the Comunidat Valencia and are responsible for JB and EF getting to know each other. This study would have not been possible without Alice Exnerová and Prof. Pavel Štys who put JB and ZB in contact. We are thankful to Centro de recuperación de fauna, Granja El Saler, for the veterinary expertise. We are thankful to José Manuel Garcia-Verdugo, his lab personnel (Patricia García-Tárraga, Arantxa Cebrian-Silla, Mariana Fill), and Mario Soriano for support in electronic microscopy techniques. We are thankful to Jan Krajíček, who conducted the initial analyses of pteridine derivatives. JB is thankful to Radek Šanda for his patience and support as Head of the Department of Zoology in the National museum. We are thankful to Jan Raška and Gerardo Antonio Cordero for comments on the manuscript. JB dedicates this study to his (growing) family.

## Ethical Statement

No human tissues were used in this study. All the turtles used in this study were captured in the wild in accordance with the Regional Decrees 14/2013 and 213/2009 on control measures on invasive alien species in the Valencian Community. All procedures performed were in accordance with the ethical standards of Direcció General de Medi Natural i d’ Avaluació Ambiental, specially the 4/1994 Regional Act on Pets Protection. All applicable international, national, and/or institutional guidelines for the care and use of animals were followed.

## Data Accessibility

Reflectance spectra data, reflectance spectra summary variables, R script of spectral analyses, reflecting platelets electromicrographs, reflecting platelets measurements and Fourier analyses output files: Dryad deposition underway, the authors will contact the journal with an updated manuscript shortly

## Authors’ Contributions

**JB** conceived the study, designed the study, coordinated the study, measured reflectance spectra, collected sample tissues, carried out the histological lab work, carried out microscopical observation, carried out the statistical analyses, carried out reflective platelets measurements and analyses, carried out Fourier analyses, participated in carotenoid analyses, drafted manuscript; **JVB** carried out the field work, communicated with local authorities; **ZB** supervised and carried out the pteridine derivatives analyses, helped to draft the manuscript; **JG** participated in Fourier analyses; **MH** carried out pteridine derivatives analyses; **KK** supervised and financially supported JB, supervised statistical analyses, helped to draft the manuscript; **PM** conceived the study, carried out the carotenoid analyses, helped to draft the manuscript; **EF** conceived the study, designed the study, financially supported the study, supervised histological and microscopical techniques, coordinated the study, drafted manuscript. All authors gave final approval for publication.

