## Supplementary materials for "Body coloration and mechanisms of colour production in Archelosauria: The case of deirocheline turtles"

**Table S1** Elution gradient of UPLC separation.

| Time [min] | 0 | 12 | 30 | 33 | 43 | 45 | 47 | 57 |
|---|---|---|---|---|---|---|---|---|
| **A** (MeCN) [%] | 20 | 100 | 5 | 0 | 0 | 100 | 20 | 20 |
| **B** (MeOH/H2O, 1:1 v/v/) [%] | 80 | 0 | 0 | 0 | 0 | 0 | 80 | 80 |
| **C** (TBME/MeCN/MeOH, 86/86/8 v/v/v) [%] | 0 | 0 | 95 | 100 | 100 | 0 | 0 | 0 |

MeCN – Acetonitrile; MeOH – Methanol; TBME – tert-Butyl Methyl Ether

**Table S2** HRAM Q-TOF MS conditions.

| | ESI ionization source | APCI ionization source |
|---|---|---|
| Capillary voltage | 4500 V | 4000 V |
| End Plate Offset | -500 V | -500 V |
| Charging Voltage | 2000 V | 2000 V |
| Nebulizer | 0.3 Bar | 2.5 Bar |
| Dry Heater | 250 °C | 250 °C |
| Dry Gas | 4.0 l/min | 4.0 l/min |
| Scan range | 60-1500 m/z | 60-1500 m/z |
| Corona curr | - | 4000 nA |
| APCI Heater | - | 450 °C |

**Table S3** SRM conditions used for LC-MS/MS determination of the pteridine derivatives.

| Compound | Molecular weight | Precursor ion | Product ion | Fragmentor (V) | Collision energy (V) |
|---|---|---|---|---|---|
| L-Sepiapterin | 237.2 | 238.2 | 192.1 | 120 | 15 |
| Pterin | 163.1 | 164.1 | 119.1 | 100 | 20 |
| Pterin-6-carboxylic acid | 209.2 | 208.1 | 162.1 | 100 | 15 |
| 6-Biopterin* | 237.1 | 238.1 | 178.1 | 115 | 17 |
| Isoxanthopterin* | 179.1 | 180.1 | 135.1 | 125 | 20 |
| Xanthopterin* | 179.1 | 180.1 | 135.1 | 125 | 20 |
| Leucopterin* | 195.1 | 196.1 | 140.1 | 120 | 16 |
| Erythropterin* | 265.1 | 266.1 | 220.1 | 110 | 8 |
| D-Neopterin* | 253.2 | 254.2 | 206.2 | 115 | 14 |

*For pteridine derivatives marked by asterisk the MS/MS conditions was adopted from Kozlík et al. (2013)[1]

[1] Kozlík P, Krajíček J, Kalíková K, Tesařová E, Čabala R, Exnerová A, Štys P, Bosáková Z. 2013 Hydrophilic interaction liquid chromatography with tandem mass spectrometric detection applied for analysis of pteridines in two Graphosoma species (Insecta: Heteroptera). *J. Chromatogr. B* **930**, 82–89

**Figure S1** Results of analyses of carotenoids. A) UPLC chromatograms (at 472nm) of mixture of carotenoid standards. B) UPLC chromatograms (at 472nm) resulting from carotenoid analyses of differen turtle skin samples with retention times of predominant carotenoids. Red chromatogram represents red postorbital region of *Trachemys scripta elegans*, all other regions are by different colors. Absorbance spectra C) of standard of lutein, D) of standard of zeaxanthin, E) of standard of cantaxanthin, F) of standard of astaxanthin

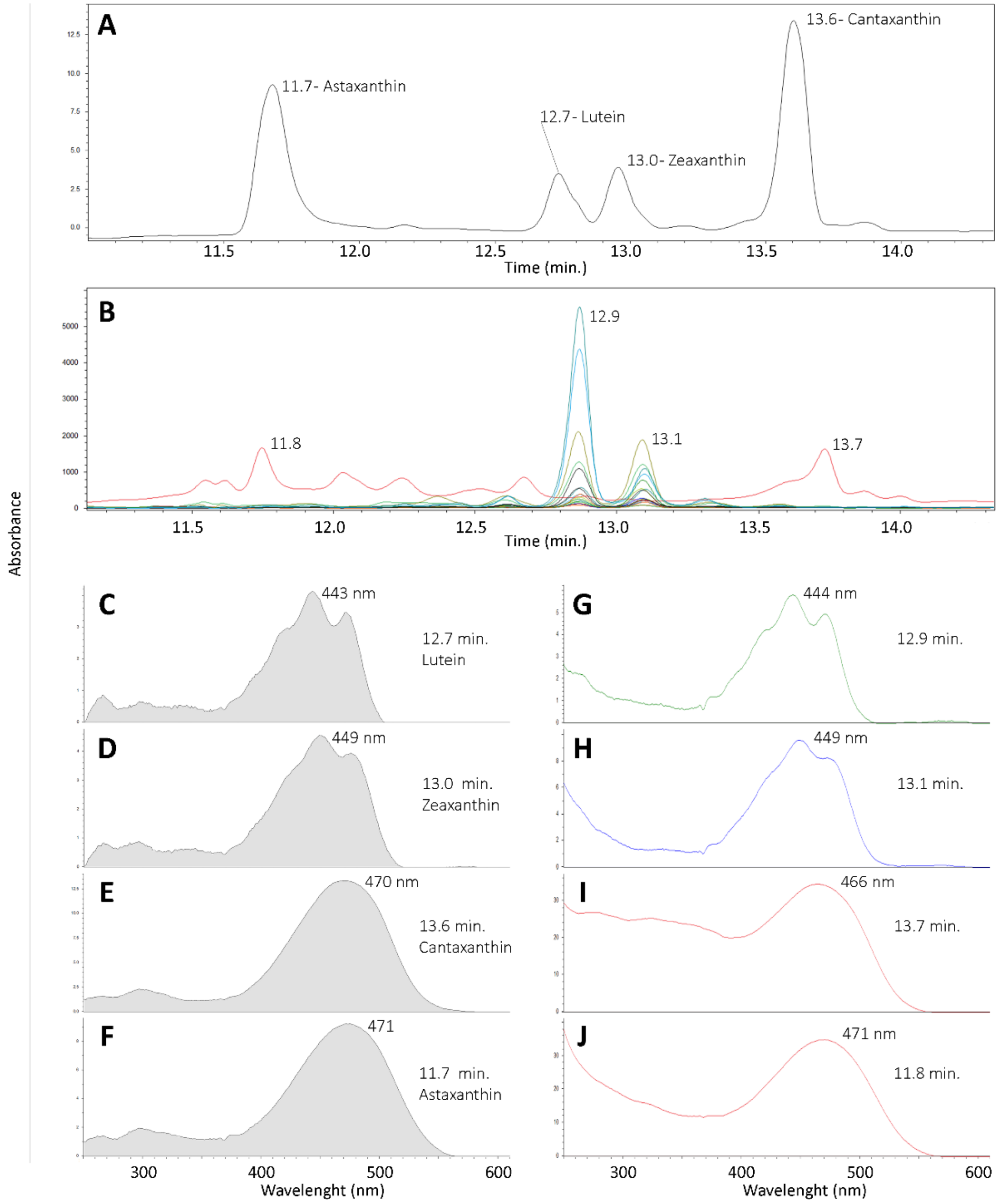

**Figure S2** Chromatograms of pteridine derivatives standards. A) TIC chromatogram of the mixture of all studied pteridine derivatives (*c* = 0.01 mg/ml), 1- L-sepiapterin, 2-pterin, 3-isoxanthopterin, 4-biopterin, 5-xantopterin, 6-leukopterin, 7-erytropterin, 8-D-neopterin, 9-pterin-6-carboxylic acid. B) SRM chromatograms of the all studied pteridine derivatives measured in the mixture (*c* = 0.01 mg/ml) under optimized LC–MS/MS conditions.

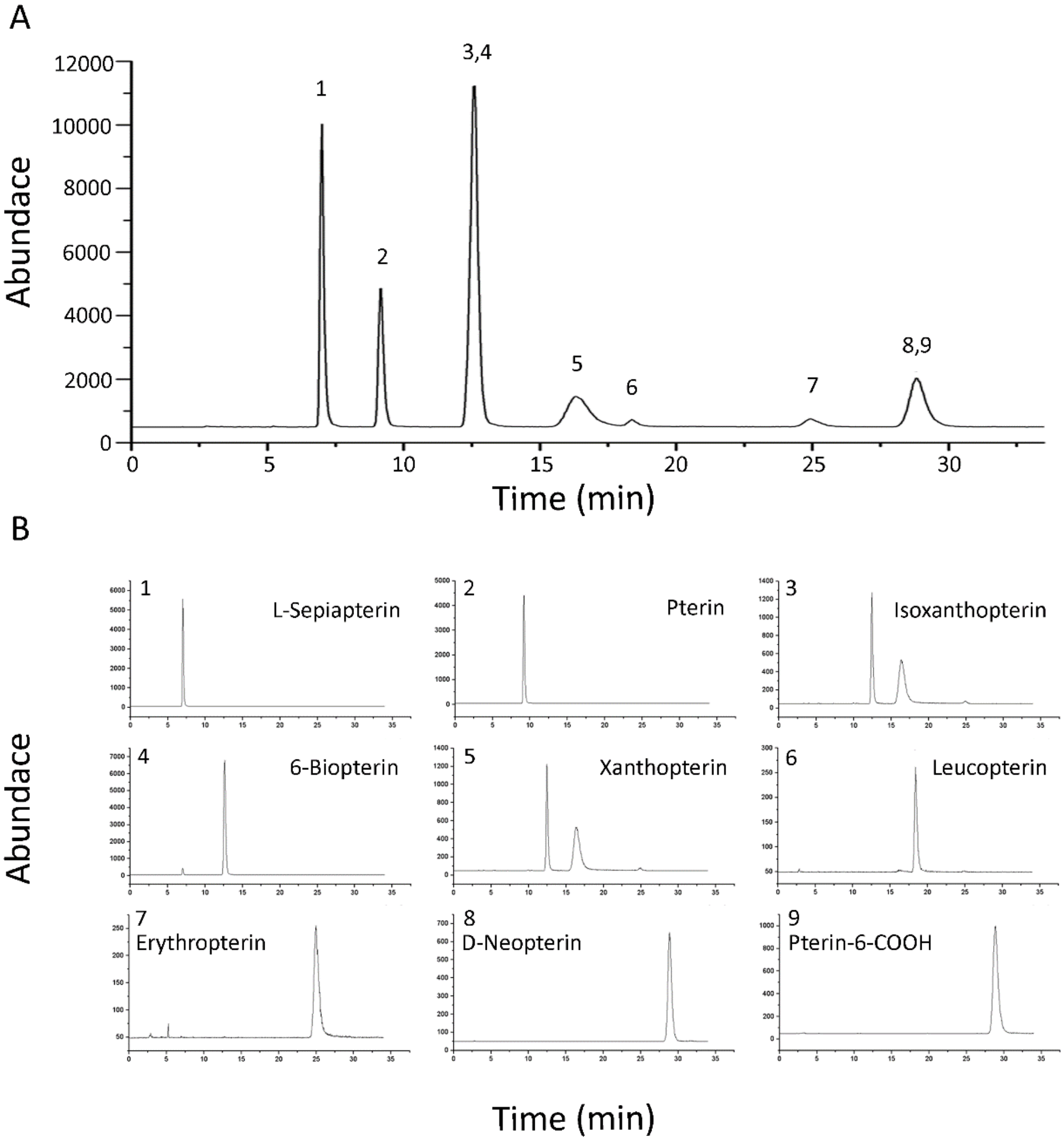

**Figure S3** Distributions of reflective platelets orientation and width. A) orientation of reflective platelets in main median chin yellow stripe (CBC) of *Pseudemys concinna*. B) orientation of reflective platelets in postorbital marking of *P. concinna*. C) orientation of reflective platelets in CBC region of *Trachemys scripta*. D) orientation of reflective platelets in main bright stripe of the fore limb (FLBS) of *T. scripta*. E) reflective platelets width of CBC region of *P. concinna*. F) reflective platelets width of PM region of *P. concinna*. G) reflective platelets width of CBC region of *T. scripta*. H) reflective paltelets width of FLBS region of *T. scripta*.

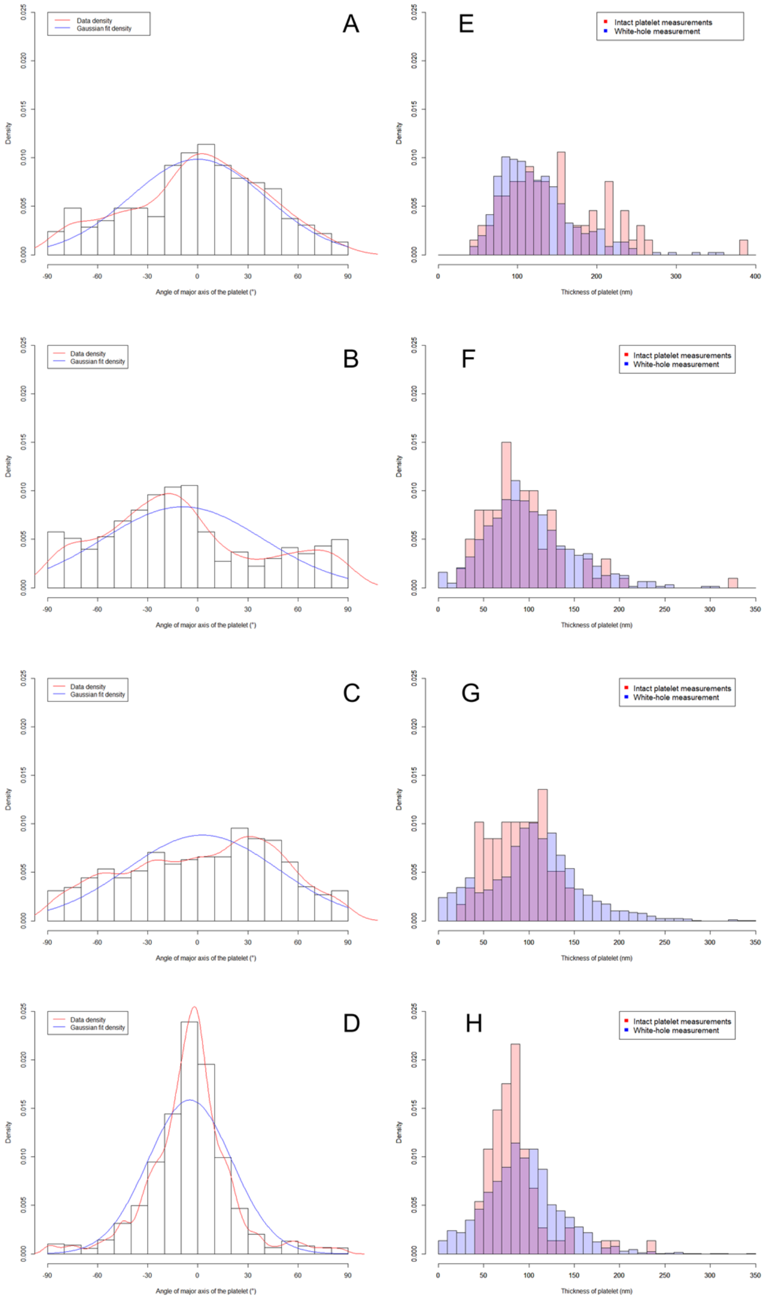

**Figure S4** Variation in 2D Fourier power spectra. A) Postorbital marking (PM) of the *Trachemys scripta*. B) Main median chin yellow stripe (CBC) of *T. scripta*. C) Main bright stripe forelimb stripe of *T. scripta*. D) PM region of *Pseudemys concinna*. E) CBC region of *P. concinna*. F) FLBS region of *P. concinna*.

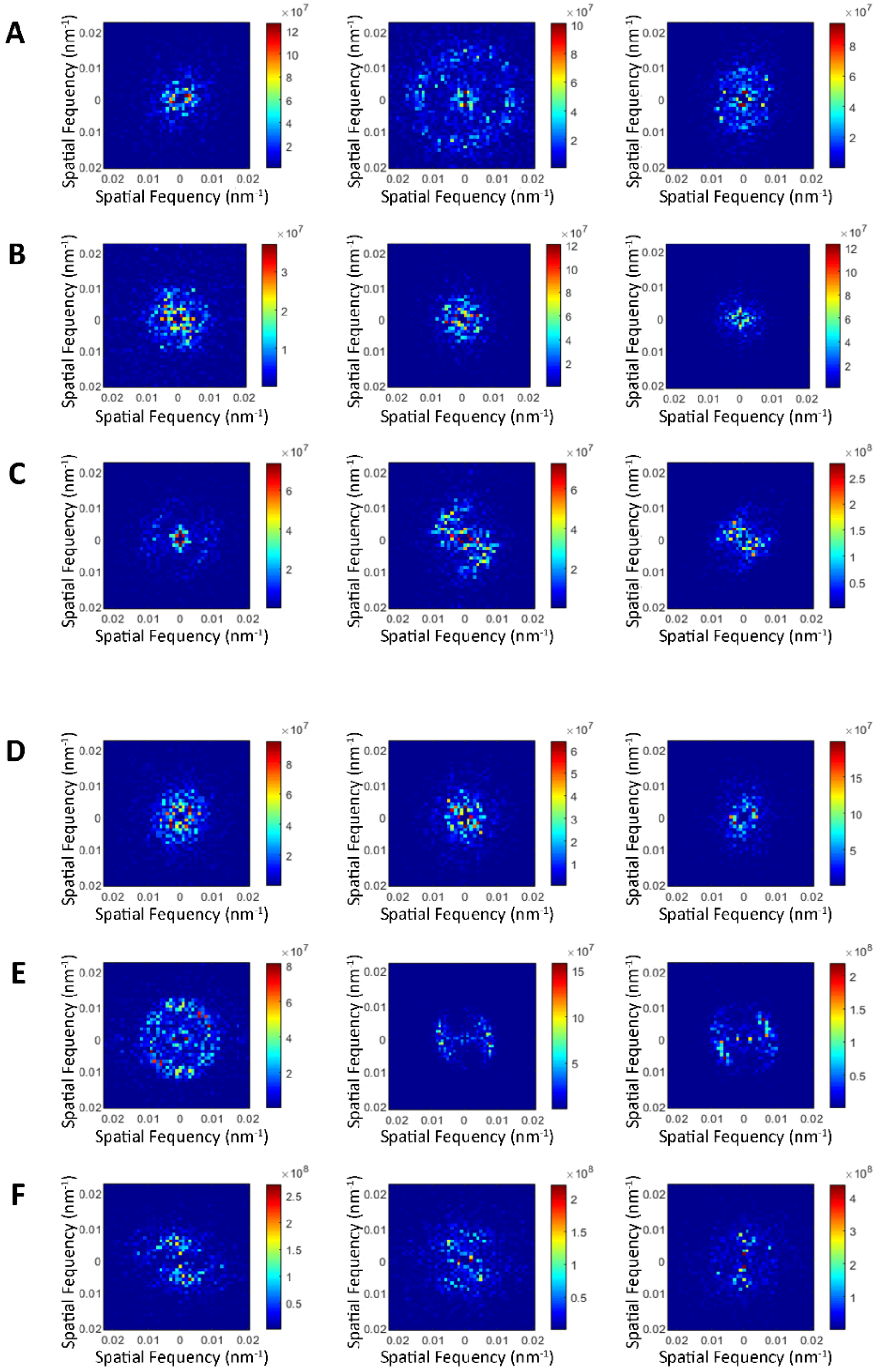

**Figure S5** Results of Fourier analyses of collagen fibers in yellow zygomatic patch YP of *Trachemys scripta scripta*. A) Radial means of Fourier power. B) Measured reflectance spectra (solid line) compared to Fourier predicted reflectivity (dashed line).

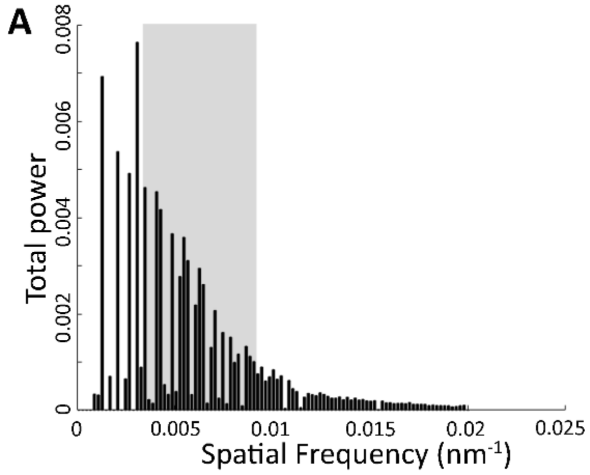

**Figure S6** Results of analyses of pteridine derivatives. A) Example of SRM chromatograms resulting from analyses of red postorbital region of *Trachemys scripta elegans*. B) Example of SRM chromatograms resulting from analyses of yellow regions of *Trachemys scripta*. C) Example of SRM chromatograms resulting from analyses of yellow regions of *Pseudemys concinna*.

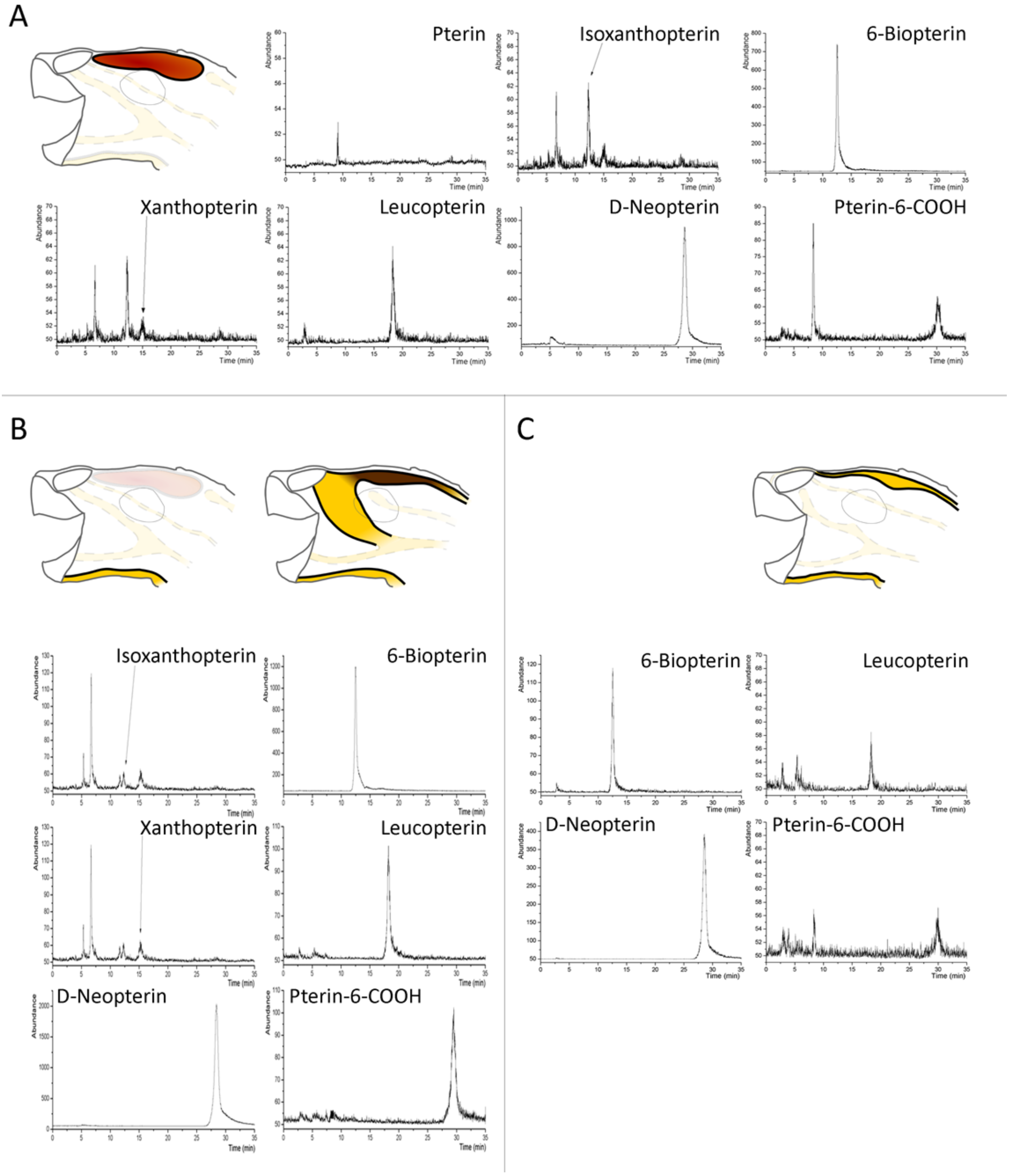

### Sexual dichromatism in *Pseudemys concinna* and *Trachemys scripta*

Previous research found some turtle species to be sexually dichromatic ( Moll et al. 1981, Bulté et al. 2013, Rowe at al. 2014)[2]. Therefore, while testing for differences among species we have controlled for the sex in the Redundancy analysis (RDA) as second constraining variable additional to species identity. RDA was performed on summary variables derived from reflectance spectra scaled to zero mean and unit variance using the package vegan in R (Oksanen et al 2015)[3] as same as it is reported in main text. Significance of results was assessed by ANOVA like permutation test for redundancy analysis (anova.cca) from vegan package with 9,999 permutations. Partition of variance of explanatory variables (species identity and sex) have been assessed using varpart function also from vegan. Further we have treated species separately. We have performed RDA again for each species, test for its significance by anova.cca and pairwise compared means of summary variables between the sexes by ANOVA with Holm's correction form multiple comparisons.

The differences between two species *P. concinna* and *T. s. elegans* as well as differences among sexes are significant (ANOVA-like permutation test; $F = 17.16$, $p < 0.001$; $F=4.66$, $p < 0.001$). Axis constrained by species identity (RDA1) explains 18 % and axis constrained by sex (RDA2) explains 4% of total variance. The first residual axis (PC1) explains 18 % and the second residual axis (PC2) explains 13 % of the total variance. However, when corrected for multiple comparison the adjusted $R^2$ for both constrained explanatory variables in the model is 0.2, while adjusted $R^2$ for model with only species identity as explanatory variable was 0.163 and for model with only sex as explanatory variable was adjusted $R^2$ 0.038.

Difference between sexes in summary variables of *P. concinna* is not significant (ANOVA-like permutation test; $F = 1.97$, $p = 0.091$). On the other hand, differences in color between sexes of *T. s. elegans* are significant (ANOVA-like permutation test; $F = 3.8954$, $p < 0.001$). Axis constrained by sex (RDA1) explains 5,5 % of total variance. First residual axis (PC1) explains 21 % and second residual axis (PC2) explains 17 % of total variance. Pairwise comparisons of summary variables between the sexes of *P.concinna* showed that there are no significant differences among sexes in neither of the summary variables (Table S4). There are significant differences in B1CBC, B1FLBS, S1.blueFLBS, and S1.redFLBS between males and females of *T. s. elegans* (Table S4).

---

[2] Moll EO, Matson KE, Krehbiel EB. 1981 Sexual and seasonal dichromatism in the Asian river turtle Callagur borneoensis. *Herpetologica* , 181–194.
Bulté G, Germain RR, O'connor CM, Blouin-Demers G. 2013 Sexual dichromatism in the northern map turtle, Graptemys geographica. *Chelonian Conserv. Biol.* **12**, 187–192.
Rowe JW, Bunce CF, Clark DL. 2014 Spectral reflectance and substrate color-induced melanization in immature and adult Midland painted turtles (Chrysemys picta marginata). *Amphibia-Reptilia* **35**, 149–159.

[3]Oksanen J *et al.* 2015 vegan: Community Ecology Package. R package version 2.0-10. 2013. *There is no Corresp. Rec. this Ref.*

**Figure S7** Differences in colour between sexes of *Pseudemys concinna* and *Trachemys scripta elegans*. A) Biplot representing results of RDA of summary variables derived from reflectance spectra of *Pseudemys concinna* and *Trachemys scripta elegans*. First axis (RDA1) is constrained by species (explains 18 % of total variance). Second axis (RDA2) is constrained by sex (explains 4 % of total variance). Dark grey hull denotes individuals of *P. concinna*, light grey hull denotes individuals of *T. scripta*, pink hull denotes females, blue hull denotes males. Summary variables are denoted by names, but when spacing would not allow to read names of variables clearly, variables are denoted by crosses. Description of summary variables are in the main text of the article. B) Biplot representing results of RDA of summary variables derived from reflectance spectra of *T. s. elegans*. First axis (RDA1) is constrained by sex (explains 5,5 % of total variance). First residual axis PC1 explains 21 % of total variance. Pink hull denotes females, blue hull denotes males. C) Results of RDA of summary variables derived from reflectance spectra of *P. concinna* with first axis constrained by sex were not significant.

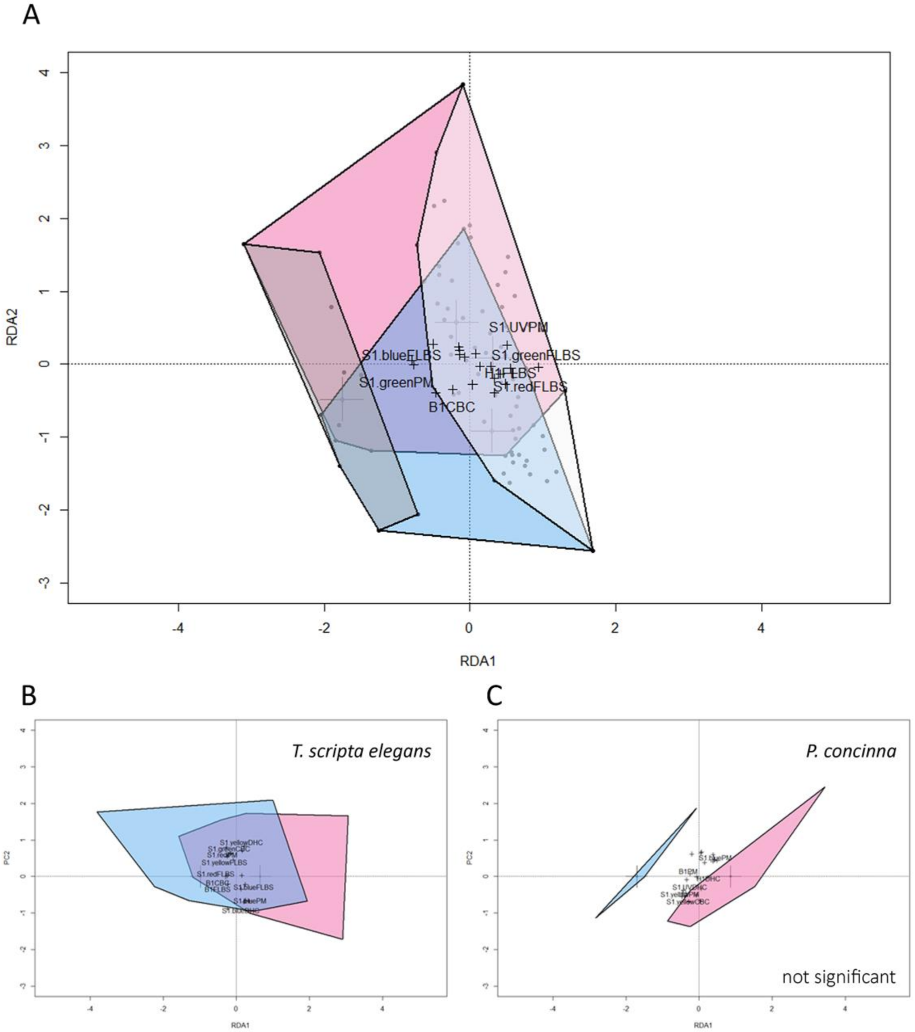

**Table S4** Results of ANOVAs of summary variables from reflectance spectra between sexes

| Species | Summary variable | F | p value | adjusted p |
|---|---|---|---|---|
| *Trachemys scripta elegans* | | | | |
| | B1CBC | 13.22 | 0.0005 | 0.0141 |
| | S1.UVCBC | 1.86 | 0.1767 | 1.0000 |
| | S1.blueCBC | 2.77 | 0.1009 | 1.0000 |
| | S1.greenCBC | 1.61 | 0.2089 | 1.0000 |
| | S1.yellowCBC | 3.42 | 0.0690 | 1.0000 |
| | S1.redCBC | 2.77 | 0.1011 | 1.0000 |
| | H1CBC | 0.67 | 0.4167 | 1.0000 |
| | B1DHC | 1.11 | 0.2949 | 1.0000 |
| | S1.UVDHC | 2.73 | 0.1035 | 1.0000 |
| | S1.blueDHC | 0.67 | 0.4158 | 1.0000 |
| | S1.greenDHC | 1.03 | 0.3149 | 1.0000 |
| | S1.yellowDHC | 0.92 | 0.3413 | 1.0000 |
| | S1.redDHC | 2.16 | 0.1466 | 1.0000 |
| | H1DHC | 1.13 | 0.2926 | 1.0000 |
| | B1FLBS | 12.98 | 0.0006 | 0.0151 |
| | S1.UVFLBS | 2.65 | 0.1084 | 1.0000 |
| | S1.blueFLBS | 10.89 | 0.0016 | 0.0375 |
| | S1.greenFLBS | 0.30 | 0.5850 | 1.0000 |
| | S1.yellowFLBS | 4.71 | 0.0336 | 0.6714 |
| | S1.redFLBS | 17.22 | 0.0001 | 0.0026 |
| | H1FLBS | 3.87 | 0.0534 | 1.0000 |
| | B1PM | 2.41 | 0.1256 | 1.0000 |
| | S1.UVPM | 5.32 | 0.0243 | 0.5096 |
| | S1.bluePM | 6.83 | 0.0111 | 0.2464 |
| | S1.greenPM | 0.96 | 0.3297 | 1.0000 |
| | S1.yellowPM | 1.92 | 0.1706 | 1.0000 |
| | S1.redPM | 6.90 | 0.0107 | 0.2464 |
| *Pseudemys concinna* | | | | |
| | B1CBC | 1.15 | 0.3086 | 1.0000 |
| | S1.UVCBC | 2.94 | 0.1169 | 1.0000 |
| | S1.blueCBC | 0.03 | 0.8741 | 1.0000 |
| | S1.greenCBC | 1.49 | 0.2495 | 1.0000 |
| | S1.yellowCBC | 1.69 | 0.2224 | 1.0000 |
| | S1.redCBC | 1.52 | 0.2458 | 1.0000 |
| | H1CBC | 0.05 | 0.8238 | 1.0000 |
| | B1DHC | 0.69 | 0.4266 | 1.0000 |
| | S1.UVDHC | 1.09 | 0.3209 | 1.0000 |
| | S1.blueDHC | 0.00 | 0.9475 | 1.0000 |
| | S1.greenDHC | 1.04 | 0.3322 | 1.0000 |
| | S1.yellowDHC | 0.38 | 0.5540 | 1.0000 |
| | S1.redDHC | 0.08 | 0.7879 | 1.0000 |
| | H1DHC | 1.20 | 0.2992 | 1.0000 |
| | B1FLBS | 0.08 | 0.7881 | 1.0000 |
| | S1.UVFLBS | 2.87 | 0.1211 | 1.0000 |
| | S1.blueFLBS | 3.69 | 0.0836 | 1.0000 |
| | S1.greenFLBS | 3.55 | 0.0890 | 1.0000 |
| | S1.yellowFLBS | 5.39 | 0.0426 | 1.0000 |
| | S1.redFLBS | 5.19 | 0.0459 | 1.0000 |
| | H1FLBS | 0.09 | 0.7726 | 1.0000 |
| | B1PM | 1.67 | 0.2259 | 1.0000 |
| | S1.UVPM | 5.87 | 0.0358 | 0.9315 |
| | S1.bluePM | 4.88 | 0.0516 | 1.0000 |
| | S1.greenPM | 2.67 | 0.1336 | 1.0000 |
| | S1.yellowPM | 8.44 | 0.0157 | 0.4239 |
| | S1.redPM | 4.27 | 0.0657 | 1.0000 |
